# G protein-coupled Receptor Contributions to Wing Growth and Morphogenesis in *Drosophila melanogaster*

**DOI:** 10.1101/2022.09.09.506847

**Authors:** Francisco J. Huizar, Nilay Kumar, Maria Unger, Vijay Velagala, Qinfeng Wu, Pavel A. Brodskiy, Jeremiah J. Zartman

**Affiliations:** Department of Chemical and Biomolecular Engineering, University of Notre Dame, Notre Dame, IN 46556, USA; Department of Bioengineering, University of Notre Dame, Notre Dame, IN, 46556, USA; Department of Biological Sciences, University of Notre Dame, Notre Dame, IN, 46556, USA

**Keywords:** G protein-coupled receptor, epithelial morphogenesis, organogenesis, developmental biology, machine learning, *Drosophila*

## Abstract

The development of multicellular organisms relies on a symphony of spatiotemporally coordinated signals that regulate gene expression. G protein-coupled receptors (GPCRs) are the largest group of transmembrane receptors that play a pivotal role in transducing extracellular signals into physiological outcomes. Emerging research has implicated neurotransmitter GPCRs, classically associated with communication in neuronal tissues, as regulators of pattern formation and morphogenesis. However, how these receptors interact amongst themselves and signaling pathways to regulate organogenesis is still poorly understood. To address this gap, we performed a systematic RNA interference (RNAi)-based screening of 111 GPCRs along with 8 G*α*, 3 G*β*, and 2 G*γ* protein subunits in *Drosophila melanogaster*. We performed a coupled, machine learning-based quantitative and qualitative analysis to identify both severe and more subtle phenotypes. Of the genes screened, 25 demonstrated at least 60% penetrance of severe phenotypes with several of the most severe phenotypes resulting from the knockdown of neuropeptide and neurotransmitter GPCRs that were not known previously to regulate epithelial morphogenesis. Phenotypes observed in positive hits mimic phenotypic manifestations of diseases caused by dysregulation of orthologous human genes. Quantitative reverse transcription polymerase chain reaction and meta-analysis of RNA expression validated positive hits. Overall, the combined qualitative and quantitative characterization of GPCRs and G proteins identifies an extensive set of GPCRs involved in regulating epithelial morphogenesis and relevant to the study of a broad range of human diseases.

## Introduction

G protein-coupled receptors (GPCRs) are the largest and most diverse group of transmembrane receptors in eukaryotic organisms (Hanlon and Andrew 2015; Nieto Gutierrez and McDonald 2018; Yang *et al*. 2021a; Sriram and Insel 2018; Insel *et al*. 2019; Tuteja 2009; Adams 2014). Due to their functional versatility, GPCRs play key roles in the regulation of transducing extracellular signals, such as peptides, proteins, and lipids, into physiological outcomes (Figure 1A) (Adams 2014; Tuteja 2009; Syrovatkina *et al*. 2016). These processes include: hormone secretion, adaptive cell immunity, cellular proliferation, metabolism, and neurotransmitter signaling (Syrovatkina *et al*. 2016; Manning *et al*. 2013; Padgett and Slesinger 2010; Ries *et al*. 2017; Schulte and Wright 2018; Schwabe *et al*. 2005; Wang 2018; Hanlon and Andrew 2015; Pal and Mukhopadhyay 2015). GPCR dysregulation has been implicated in numerous disease categories including rheumatic, neurological, pulmonary, cardiac, endocrine, and epithelial (Figure 1A) (Skiba and Kruse 2021). Because of this, GPCRs are highly attractive therapeutic targets (Yang *et al*. 2021a; Sriram and Insel 2018; Insel *et al*. 2019). Although some GPCRs have been extensively studied, newly discovered GPCRs and their functions, especially in relation to their role during early developmental stages and organogenesis, remain to be elucidated (Belgacem and Borodinsky 2011; Hanlon and Andrew 2015; Pal and Mukhopadhyay 2015; Schulte and Wright 2018).

**Figure 1.**
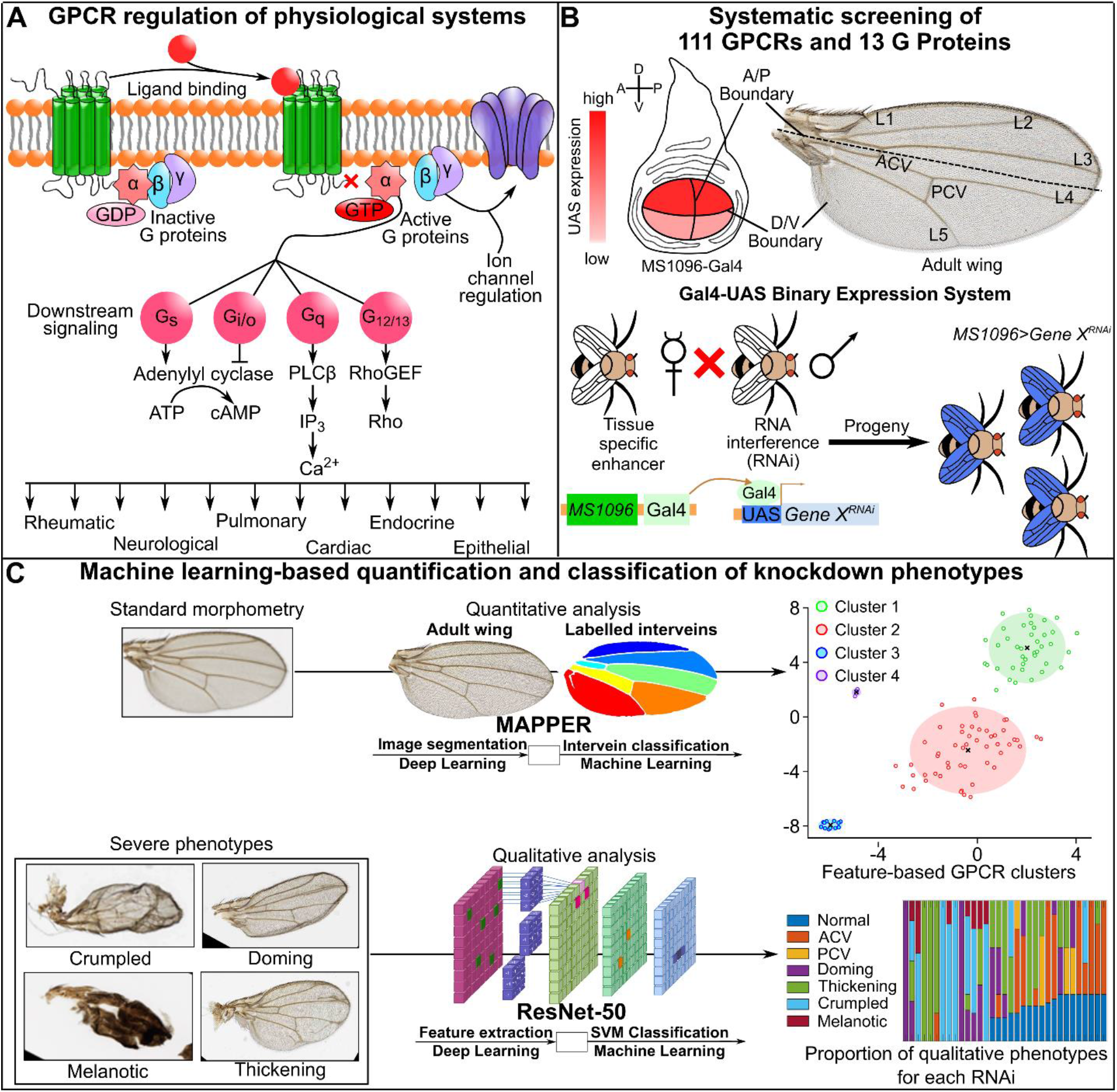
Machine learning-based high-throughput screening of GPCRs in *Drosophila* wing morphogenesis. **(A)** G protein-coupled receptors (GPCRs) are cell surface sensors that upon receiving an input signal, such as a ligand binding to the receptor, begin a downstream signaling cascade through G protein subunits G*α*, *Gβ*, and G*γ*. Upon ligand binding, conformational changes induce G protein subunit activation as guanosine triphosphate (GTP) replaces bound guanosine diphosphate (GDP). Depending on the activated G*α* sub-type (G_*s*_, G_*i/o*_, G_*q*_, or G_12/13_), various downstream signaling molecules are recruited to regulate physiological processes. The dissociated G*βγ* subunit can then act as an ion channel regulator. **(B)** We used the Gal4-UAS expression system to silence gene expression via RNA interference (RNAi) of a desired gene of interest (UAS-Gene *X^RNAi^*) in the *Drosophila* wing. The MS1096-Gal4 driver has a higher expression level in the dorsal compartment of the developing *Drosophila* wing disc pouch than in the ventral compartment. The adult wing has distinct morphological features: longitudinal veins (L1-L5), the anterior crossvein (ACV), and the posterior crossvein (PCV) that we rely on to study the effect of GPCRs on morphogenesis. A systematic knockdown screen of 111 GPCRs and 13 G proteins was carried out. A: anterior, P: posterior, D: dorsal, V: ventral. **(C)** Combinations of machine learning and deep learning were used for high-throughput screening and analysis of adult wing images. Wings with standard morphometry underwent quantitative analysis, while wings with severe phenotypes underwent qualitative analysis. Coupled analyses enable a more comprehensive characterization of the effects of GPCR on epithelial morphogenesis.

One of such cases of unknown GPCR functionality is the role of neuropeptide and neurotransmitter GPCRs in the patterning and development of multicellular organisms. Development of multicellular organisms relies on a collection of spatiotemporally coordinated signals to drive gene expression of regulators of cellular proliferation, differentiation, cell-cell communication, and motility (Blau and Baltimore 1991; Adams and Watt 1993). The *γ*-aminobutyric acid (GABA) and 5-hydroxytryptamine (5-HT) families of GPCRs are well-documented in their roles for modulating neural activity, however, their contribution to morphogenesis of organs outside of the nervous system is still incompletely characterized (Hannon and Hoyer 2008; Terunuma 2018; Barnes and Sharp 1999; Bettler *et al*. 2004; Pinard *et al*. 2010).

GPCRs can be categorized into six classes based on their amino acid sequences and functional similarities: Class A (rhodopsin-like family), Class B (secretin and adhesion family), Class C (metabotropic glutamate receptors), Class D (fungal mating pheromone receptors), Class E (cyclic adenosine monophosphate (cAMP) receptors), and Class F (Frizzled and Smoothened receptors) (Ghosh *et al*. 2015; Foord *et al*. 2005; Lee *et al*. 2018; Yang *et al*. 2021a). Beyond the estimated 800 known human GPCRs, there exist many receptors in the human genome that have amino acid sequences similar to known GPCRs with unknown activating ligands and signaling mechanisms (Foord *et al*. 2005). Thus, further characterization of these probable orphan GPCRs, their functionalities, and their corresponding classes are a focal point for expanding the list of therapeutic targets for many diseases.

*Drosophila melanogaster* is a classic model organism for studying human diseases and organ development (Mirth and Shingleton 2012; Mirzoyan *et al*. 2019; Pandey and Nichols 2011; Jennings 2011). GPCRs are the largest group of receptors in *Drosophila* with an estimated 111 GPCRs that signal through a combination of 8 G*γ*, 3 G*β*, and 2 G*γ* subunits (Hanlon and Andrew 2015; Thurmond *et al*. 2019). G protein subunits signal primarily by regulation of the dynamics of various second messengers, such as cAMP and calcium ions (Ca^2+^), wherein second messenger signaling impacts downstream cell function (Figure 1A) (Brodskiy *et al*. 2019; Hanlon and Andrew 2015; Hepler and Gilman 1992; Khan *et al*. 2013). Many neurotransmitter-related GPCRs have been thoroughly studied in adult *Drosophila* and have classical roles in adult behavior (Hanlon and Andrew 2015; Manning *et al*. 2013; Schwabe *et al*. 2005). However, while many of these are expressed during embryogenesis, considerably less is known about their roles in development. With increasing evidence demonstrating multiple GPCRs are vital to the development of the *Drosophila* wing (Sobala and Adler 2016; Ren *et al*. 2005), a premier model of epithelial morphogenesis, there has not been a concerted effort to comprehensively evaluate which of the 111 GPCRs have roles in *Drosophila* wing development (Fristrom 1988; Etournay *et al*. 2015; Ayers and Thérond 2010; Schulte and Wright 2018; Blair 2007; Garcia De Las Bayonas *et al*. 2019).

*Drosophila* provides many advantages for identifying and characterizing conserved components of signal transduction pathways. These include fast life cycle, abundance of available genetic tools, highly conserved homology to the human genome, and cheap husbandry (Ashburner *et al*. 2005; Jennings 2011; Perrimon *et al*. 2016; Pandey and Nichols 2011). These advantages enable rapid phenotypic screening of genes in *Drosophila* with results that are directly relevant to human biology (Belacortu and Paricio 2011; Perrimon *et al*. 2016; Kumar *et al*. 2022). In particular, the *Drosophila* wing has served as a viable model system for identifying and studying genes and biophysical mechanisms important for receptor crosstalk, growth, pattern formation, and morphogenesis (López-Varea *et al*. 2021a,b; Rotelli *et al*. 2019; George *et al*. 2019; Saad and Hipfner 2021; Heigwer *et al*. 2018). The *Drosophila* wing exhibits distinct morphological characteristics, including seven intervein regions, five longitudinal veins, two crossveins, and trichomes -the hair-like structures along the surface and edge of the wing (Figure 1B). Quantitative and qualitative changes in these morphological features can largely provide insight into the underlying biological mechanisms that regulate the development of the wing (Buchmann *et al*. 2014; Narciso and Zartman 2018; Restrepo *et al*. 2014; Strigini and Cohen 1999; Kumar *et al*. 2022). More specifically, targeted genetic perturbations induced by RNA interference (RNAi) can be used to uncover novel biological insights and future research directions in developmental biology using *Drosophila* (Kumar *et al*. 2022; George *et al*. 2019; Saad and Hipfner 2021).

Here, we report a systematic RNAi-based investigation into the phenotypes associated with inhibition of 111 GPCRs along with 8 G*α*, 3 G*β*, and 2 G*γ* proteins during *Drosophila* wing development (Figure 1B). For quantitative analysis, we employed our comprehensive pipeline, MAPPER (Kumar *et al*. 2022), to perform high-content genetic wing screening via deep learning for image segmentation and machine learning for feature classification (Figure 1C). For qualitative analysis, the ResNet-50 convolutional neural network (He *et al*. 2015; LeCun *et al*. 2015) was used in tandem with a support-vector machine (Wang 2005) for classification of severe *Drosophila* wing phenotypes (Figure 1C, SI Figures 1 and 2). With the coupled analysis, we discovered several classes of GPCRs that demonstrate severe phenotypic irregularities when knocked down. Of the GPCRs and G proteins screened, 25 demonstrated at least 60% penetrance of severe qualitative phenotypes with 12 neuropeptide receptors that resulted in a change of at least 10% of the total area of the wing compared to the control group. Interestingly, several of the most severe phenotypes were the result of RNAi of neuropeptide and neurotransmitter receptors, many of which are reported to have low-to-no expression in the developing *Drosophila* wing disc (Ren *et al*. 2005; Sobala and Adler 2016; Celniker *et al*. 2009). Quantitative reverse transcription polymerase chain reaction (RT-qPCR) and cross-comparison with other RNA-seq studies confirmed expression of candidate hits in wing disc cells. The results of the RT-qPCR experiments suggest that even low abundance GPCRs may lead to severe phenotypic outcomes when dysregulated. Of note, the observed phenotypes of positive hits serve as phenologs (phenotypic manifestations) of orthologous human genes that are implicated in a broad range of diseases (Table 1).

**Table 1.** Human diseases associated with high similarity *Drosophila* orthologs identified in knockdown screen.

| *Drosophila* gene | BL# screened<sup>a</sup> | Observed phenotype | GPCR Class | Human ortholog<sup>b</sup> | Associated diseases | Disease phenotype |
| --- | --- | --- | --- | --- | --- | --- |
| *Gβ5* | 28310 | Doming, PCV defect | G protein | *GNB5* (83%) | Intellectual developmental disorder with cardiac arrhythmia (IDDCA) (De Nittis et al. 2021) | Early-onset intellectual disability, bradycardia, and cardiac arrest (Shao et al. 2021) |
| *Gαs* | 50704 | Melanotic | G protein | *GNAL* (81%) | Dystonia (Deutschländer and Wszolek 1993) | Dystonic tremors (Carecchio et al. 2016) |
| *Gγ30A* | 34484 | Doming, Melanotic | G protein | *GNG13* (64%) | Breast cancer (Liu et al. 2022b) | Malignant cancer growth (Liu et al. 2022b) |
| *TkR99D* | 55732 | Lethal | Class A | *TACR3* (63%) | Oral squamous cell carcinoma (Obata et al. 2016) | Tumorigenesis and mandibular bone destruction (Obata et al. 2017) |
| *fz* | 34321 | Thickening, Vein bifurcation | Class F | *FZD7* (61%) | Various types of cancer (Michelli et al. 2020) | Organ metastasis and poor clinical prognosis (Zhang and Xu 2022) |
| *CCHa1-R* | 51168 | Thickening, Vein bifurcation | Class A | *GRPR* (59%) | Various types of cancer (Morgat et al. 2017) | Increased cancer cell migration (Patel et al. 2014) |
| *CapaR* | 27275 | Doming, ACV defect | Class A | *NMUR2* (56%) | Breast cancer (Garczyk et al. 2017) | Tumorigenesis (Lin et al. 2015) |
| *mtt* | 44076 | Melanotic | Class C | *GRM1* (52%) | Triple-negative breast cancer (Bastiaansen et al. 2020) | Mediation of tumor cell growth and endothelial cell proliferation (Sexton et al. 2018) |
| *Rh2* | 77340 | Thickening, ACV defect | Class A | *OPN4* (52%) | Melanoma (de Assis et al. 2022) | Knockout results in faster cell cycle progression (de Assis et al. 2021) |
| *5-HT1B* | 54006, 27632 | Crumpled, Thickening | Class A | *HTR1A* (45%) | Breast and lung cancer (Kopparapu et al. 2013) | Knockdown increases metastasis (Liu et al. 2022a) |
| *CG30340* | 28652 | Doming | Class A | *GPR3* (41%) | Alzheimer's Disease (Huang et al. 2015) | Loss of GPR3 improves memory in Alzheimer's mouse model (Huang et al. 2015) |

<sup>a</sup> BL: Bloomington *Drosophila* Stock Center<sup>b</sup> Amino acid similarity score % obtained from DRSC Integrative Ortholog Prediction Tool (DIOPT) version 8.0 (Hu et al. 2011). Human orthologs displayed have the best score and best reverse score reported.

Interestingly, a quantitative comparison of phenotypic similarities between positive hits using Gaussian mixture models and Euclidean distances of dimension reduced wing features (Yang *et al*. 2012) revealed the identification of multiple phenotypic clusters. This cluster-based analysis leads to a prediction of novel protein-protein interactions between GPCRs, including for the *Drosophila* 5-HT1B receptor (ortholog of human *HTR1A*), which produced one of the most severe phenotypes. The predicted protein-protein interactions for the 5-HT1B receptor are supported by comparison with the STRING protein-protein interaction network database (Szklarczyk *et al*. 2015, 2011). Unconfirmed protein-protein interactions suggest new biological insights and avenues to uncover the clinical implication of serotonin receptors with other GPCRs in a range of neurological conditions, including depression, bipolar disorder, schizophrenia, Alzheimer’s disease, and cognitive function (López-Figueroa *et al*. 2004; Yang *et al*. 2021b; Tiger *et al*. 2018; Garcia-Alloza *et al*. 2004). Overall, the combined machine learning approaches for both qualitative and quantitative analyses enables a more comprehensive characterization of new regulators of epithelial morphogenesis. Overall, the results of this screen provide a starting point for further exploration of the molecular mechanisms underlying GPCR regulation and cross-talk during epithelial morphogenesis.

## Materials and methods

### Identification of GPCR screening library

The list of 111 GPCRs was obtained from the FlyBase Gene Group titled: “Gene Group: G PROTEIN COUPLED RECEPTORS.” This was accessed using FlyBase ID FBgg0000172 of FlyBase version: FB2019_01 (Thurmond *et al*. 2019). The list of 13 G proteins was obtained from the FlyBase Gene Group titled: “Gene Group: HETEROTRIMERIC G-PROTEIN SUBUNITS.” This was accessed using FlyBase ID FBgg0000458 of FlyBase version: FB2019_01 (Thurmond *et al*. 2019). Only *Drosophila* strains that had readily available stocks from the Bloomington *Drosophila* Stock Center were screened. Supplementary File S1 provides the comprehensive list of genes and their associated Bloomington *Drosophila* Stock Center number. Supplementary File S2 includes the list of the screened genes, their FlyBase IDs, their GPCR class, and general gene information.

### *Drosophila* stocks and culture

*Drosophila melanogaster* strains were obtained from the Bloomington *Drosophila* Stock Center as indicated by stock number (BL#). RNAi lines selected were those generated by the Transgenic RNAi project from the functional genomics platform at Harvard Medical School (Perkins *et al*. 2015). When possible, multiple, independent RNAi lines were tested for each gene investigated. *Drosophila* were raised at 25 °C and 12-hour light cycle on the standard Bloomington *Drosophila* Stock Center cornmeal food recipe.

### *Drosophila* genetic crosses

The Gal4-UAS binary expression system was utilized to express the RNAi constructs for the identified genes in the developing *Drosophila melanogaster* wing (Duffy 2002). The MS1096-Gal4 (BL#25706) line was used as the basis for genetic crosses, which drives gene expression in the developing *Drosophila* wing disc with more pronounced expression in the dorsal compartment (Figure 1B) (Lindström *et al*. 2017; Neumann and Cohen 1996; Lin and Goodman 1994; Capdevila and Guerrero 1994). There is conflicting evidence of either no expression or weak expression patterns in the central nervous system (CNS) when using the MS1096-Gal4 driver (*Lindström et al*. 2017;Ray and Lakhotia 2019). However, to achieve a more efficient screening of GPCRs and G proteins, we focused on observation of wing phenotypes only.

Genetic knockdown progeny were generated by crossing the MS1096-Gal4 line to RNAi-based transgenic lines (UAS-Gene *X^RNAi^* (Perkins *et al*. 2015)). The RNAi for the ryanodine receptor (RyR) (BL#31540) was used as a background control for MS1096-Gal4>UAS-RNAi crosses. We have previously demonstrated the MS1096-Gal4 × UAS-RyR^*RNAi*^ cross does not exhibit significant morphological or size defects when compared to wild type controls due to the RyR gene not being expressed in the *Drosophila* wing disc (Brodskiy *et al*. 2019; Gramates *et al*. 2017). Thus, MS1096-Gal4 × UAS-RyR*RNAi* progeny enable assessment of the impact of knocking down GPCRs and G proteins in the wing disc during development. F1 progeny from MS1096-Gal4>UAS-RNAi crosses were compared to those of the F1 progeny of the MS1096-Gal4 × UAS-RyR^*RNAi*^ cross. Only wings from male F1 progeny emerging from the crosses were scored to avoid data variation due to sex. When possible, multiple crosses were generated for each RNAi line for additional biological replicates until approximately 15 samples per cross were available.

Heterozygous MS1096-Gal4 expressing flies contain venation defects with variable penetrance (George *et al*. 2019). Less than 5% of MS1096>UAS-RyR^*RNAi*^ F1 progeny had venation defects or severe phenotype penetrance with a 95% confidence interval of 0.845% - 22.7% using a one-sample proportions test without continuity correction. Only MS1096>UAS-RNAi progeny populations with at least 40% venation penetrance or at least 60% penetrance of severe phenotypes were considered in the qualitative analysis. These thresholds were set to reduce the likelihood of including false positives among the candidates of interest. In the case of multiple independent RNAi lines being available to examine, we report positive/negative hits with respect to the specific BL# and acknowledge this may introduce bias to genes with only one RNAi line available.

Virgin MS1096-Gal4 females were crossed with males from each UAS-RNAi strain. Female virgins were collected prior to eclosure by confirming the absence of sex combs in the pupal casing. Approximately nine female MS1096-Gal4 virgins were crossed with five male UAS-RNAi flies. Wings were mounted on glass microscopy slides. One wing was extracted from each fly, placed in ethanol, and approximately 15 wings per cross were mounted on each slide in 40 *μ*L Permount medium (Fisher Scientific, SP15). Because crossing procedures were standardized for each cross, variance in samples across vials was not investigated to conduct a more efficient screen. A glass coverslip was placed on top of wings embedded in the Permount medium to mount the samples for long-term storage. A small weight was added to the top of the coverslip to evenly distribute the Permount medium and flatten the wings. The slides were then labeled using a QR code system (Dymo LabelWriter 450).

### High-throughput imaging and image processing

Slides were batch imaged using an Aperio slide scanner (Leica BioSystems) at 5X magnification (Courtesy of South Bend Medical Foundation). Slides were stored coverslip-side up at room temperature. Image data was stored using the University of Notre Dame Center for Research Computing. The resulting SVS files from the slide scanner were processed using the pixel classification platform Ilastik (Sommer *et al*. 2011) to generate segmentation masks of the wings. A MATLAB script from our wing image analysis pipeline (Kumar *et al*. 2022) was used to crop individual wings from the SVS files for further analysis.

### Reverse transcription-quantitative polymerase chain reaction of wing disc cells

RT-qPCR was performed on cultured third instar larval wing disc cells (CME W1 Cl.8+, *Drosophila* Genomics Resource Center Stock 151) (Peel and Milner 1990). Cells were cultured using the recommended Shields and Sang M3 insect medium with 2% fetal bovine serum, 5 *μ*g/mL insulin, and 2.5% fly extract. Fly extract was prepared using the available protocol provided by the *Drosophila* Genomics Resource Center (Cherbas 2016). Cells were grown in T25 flasks and maintained between 2.0 × 10^6^ and 1.0 × 10^7^ cells/mL. Cells were incubated at 25 °C in a standard incubator without CO_2_ exchange (Luhur *et al*. 2019).

TRIzol^*™*^ reagent (catalog #15596026, Invitrogen, ThermoFisher Scientific) and the corresponding user manual (Pub. No. MAN0001271) was used to extract mRNA from *Drosophila* Cl.8 cells. KiQcStart SYBR Green Primers (MilliporeSigma catalog #KSPQ12012G) were used for RT-qPCR reactions. Gene targets of KiQcStart primers, their corresponding labels throughout the text, their FlyBase ID, and their NCBI Reference Sequence ID are found in Table 2. *α*-Tubulin at 84B (NCBI Reference Sequence: NM_057424), a ubiquitously expressed gene, was used as a positive control. No template control (NTC) wells containing no template DNA were used as the negative control. For select gene targets, multiple oligonucleotide primers were evaluated, and they are denoted as distinct numbers in parentheses. RT-qPCR reactions were carried out using the Quanta Biosciences*^™^* one-step SYBR Green RT-qPCR kit (catalog #95087) and the Applied Biosystems StepOne^*™*^ Real-Time PCR System (ThermoFisher Scientific). Reactions were carried out in triplicate.

**Table 2.** MilliporeSigma KiQcStart SYBR Green Primers used for RT-qPCR experiments.

| Manuscript label | Gene Target | Primer Pair ID<sup>a</sup> | FlyBase ID | NCBI RefSeq ID |
| --- | --- | --- | --- | --- |
| α-Tubulin | α-Tubulin at 84B | DMEL_alphaTub84B_1 | FBgn0003884 | NM_057424 |
| Negative | No template primers | None | None | None |
| Gαf (1) | G protein $\alpha$ f subunit | DMEL_Galphaf_1 | FBgn0010223 | NM_079394 |
| Gαf (2) | G protein $\alpha$ f subunit | DMEL_Galphaf_2 | FBgn0010223 | NM_079394 |
| Gαs (1) | G protein $\alpha$ s subunit | DMEL_Galphas_1 | FBgn0001123 | NM_058158 |
| Gαs (2) | G protein $\alpha$ s subunit | DMEL_Galphas_2 | FBgn0001123 | NM_058158 |
| Gβ13F | G protein $\beta$ subunit 13F | DMEL_Gbeta13F_1 | FBgn0001105 | NM_080351 |
| Gγ30A | G protein $\gamma$ subunit 30A | DMEL_Ggamma30A_13 | FBgn0267252 | NM_080068 |
| CCHa1R (1) | CCHamide-1 receptor | DMEL_CCHa1-R_2 | FBgn0050106 | NM_137397 |
| CCHa1R (2) | CCHamide-1 receptor | DMEL_CCHa1-R_3 | FBgn0050106 | NM_137397 |
| HT1B (1) | 5-HT receptor 1B | DMEL_5-HT1B_1 | FBgn0263116 | NM_001169730 |
| HT1B (2) | 5-HT receptor 1B | DMEL_5-HT1B_2 | FBgn0263116 | NM_001169730 |
| mthl3 (1) | methuselah-like 3 | DMEL_mthl3_1 | FBgn0028956 | NM_145335 |
| mthl3 (2) | methuselah-like 3 | DMEL_mthl3_2 | FBgn0028956 | NM_145335 |
| CG13579 | CG13579 | DMEL_CG13579_1 | FBgn0035010 | NM_138073 |
| CG30340 | CG30340 | DMEL_CG30340_1 | FBgn0050340 | NM_165686 |
| GABA-B-R1 | GABA-B receptor 1 | DMEL_GABA-B-R1_1 | FBgn0260446 | NM_078845 |
| GABA-B-R2 | GABA-B receptor 2 | DMEL_GABA-B-R2_1 | FBgn0027575 | NM_079714 |
| HT2A | 5-HT receptor 2A | DMEL_5-HT2A_1 | FBgn0087012 | NM_001170035 |
| mtt | mangetout | DMEL_mtt_10 | FBgn0050361 | NM_206058 |

<sup>a</sup> Primer pair IDs can be found by entering the NCBI RefSeq ID into the KiQcStart<sup>TM</sup> Primers web page search tool

The quantification of amplification is reported as the normalized fluorescent signal of the reporter dye in a sample (R*n*). The difference in normalized fluorescent signal from the experimental reaction and the baseline signal generated by the StepOne*^™^* Software are denoted as Δ*Rn*, which provides a measure of amplification over time throughout the experiment (Figure 4A-C). Quantification can be further compared by observing the cycle threshold (C*T*), which is the fractional cycle number at which the fluorescent signal passes the threshold determined by the StepOne*^™^* Software. C*T* levels are thus inversely proportional to the amount of target cDNA in the sample (*i.e*., the lower the measured C*T* level, the greater the amount of target cDNA is present in the sample). Δ*CT* is a measure that demonstrates differences in expression between a target gene of interest and a ubiquitously expressed positive control gene (*α*-Tubulin at 84B), by subtracting the C*T* of a gene of interest from the C*T* of the positive control (Figure 4A’-C’).

**Figure 4.**
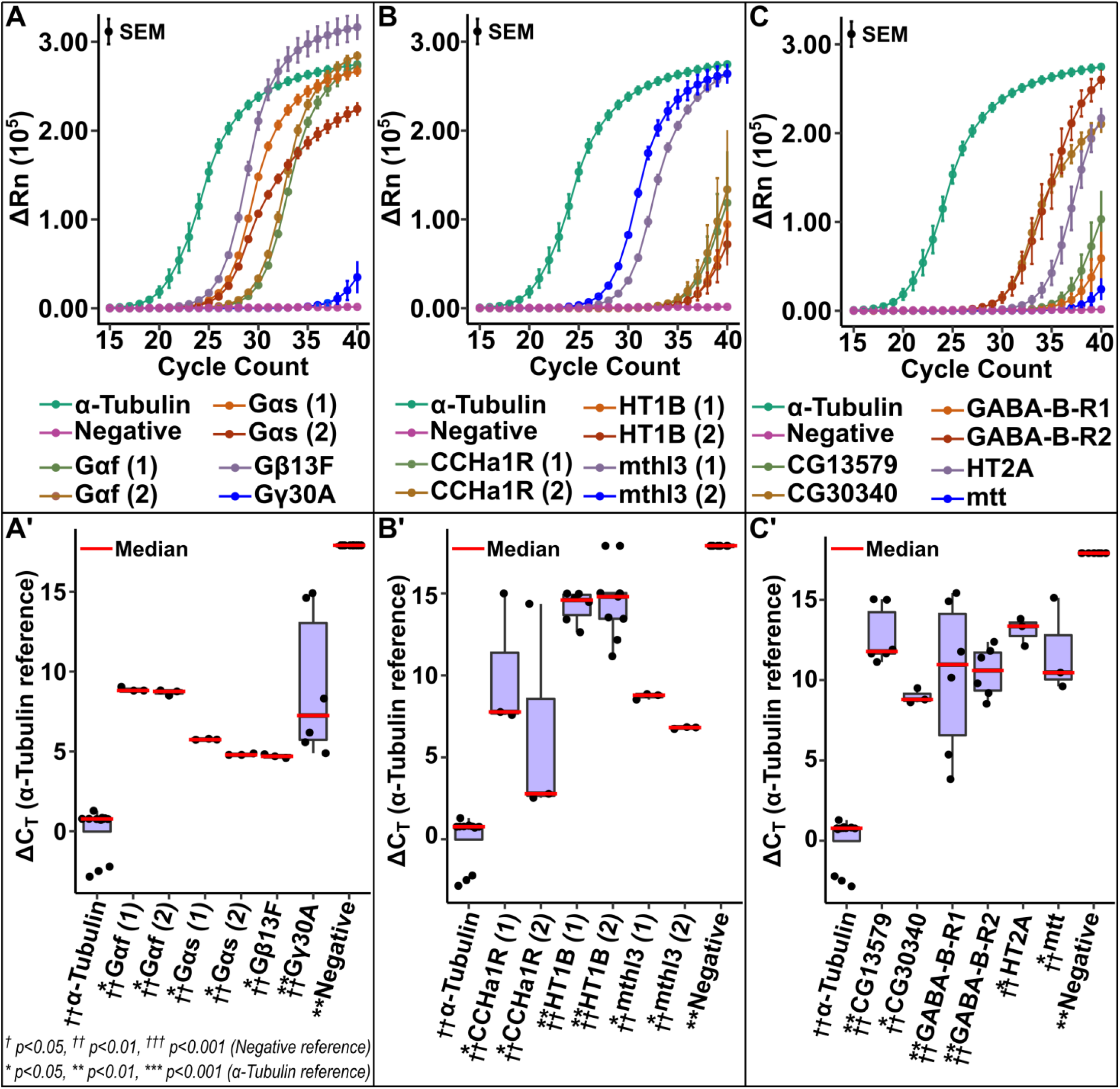
Reverse transcription-quantitative polymerase chain reaction (RT-qPCR) confirms expression of positive hits in *Drosophila* wing imaginal disc. RT-qPCR was performed on cultured third instar larval wing disc cells (CME W1 Cl.8+, DGRC Stock 151). Select positive hit genes that exhibited severe wing phenotypes were tested to confirm expression in the *Drosophila* imaginal wing disc. *α*-Tubulin at 84B (NCBI Reference Sequence: NM_057424), a ubiquitously expressed gene, was used as a positive control. No template control (NTC) wells containing no template DNA were used as a negative control. **(A-C)** The difference in normalized fluorescent signal from the experimental reaction and the baseline signal generated by the StepOne^*™*^ Software are plotted (Δ*R_n_*). Error bars represent the standard error of the mean (SEM) of each gene at a given cycle count. **(A’-C’)** The difference in the number of amplification cycles required to reach the StepOne^*™*^ Software generated threshold between the experimental gene and the positive control are plotted (Δ*C_T_*). Experimental runs in which no amplification was detected were given *C_T_* values of 40 because that was the extent of the experimental runs. Pairwise Mann-Whitney U tests were performed for statistical analysis with false discovery rate (FDR) correction (* *p* < 0.05, ** *p* < 0.01, *** *p* < 0.001 compared to positive control; ^†^ *p* < 0.05, ^††^ *p* < 0.01, ^†††^ *p* < 0.001 compared to negative control). **(A-A’)** Δ*R_n_* and Δ*C_T_* plots for various G proteins. Numbers in parentheses indicate use of multiple, distinct oligonucleotide primers. **(B-B’)** Δ*R_n_* and Δ*C_T_* plots for various GPCRs where multiple, distinct (numbers in parentheses) oligonucleotide primers were evaluated. **(C-C’)** Δ*R_n_* and Δ*C_T_* plots for various GPCRs in which only one pair of oligonucleotide primers was evaluated.

### Training the qualitative phenotype classifier

The image screening pipeline required processing of over several thousand images, analysis of which becomes intractable manually. Thus, a robust, automated algorithm that can precisely measure and extract measurements from wing images while simultaneously handling all edge cases is highly desired. To ensure robust quantification and qualitative analyses of the data set, the first step of the pipeline aimed to classify raw wing images into different classes of interest: Crumpled, Doming, Melanotic, Normal, or Thickening (SI Figures 1 and 2). In addition to detecting severe phenotypes that resulted from genetic perturbations, this classification step also ensured the removal of images that had poor lens focus, torn samples, and mounting artifacts. A similar image filtering method was used in our previously reported open-source pipeline for high-throughput image processing and measurement of *Drosophila* wings (Kumar *et al*. 2022).

Qualitative features of *Drosophila* wing data for wings whose quantitative measures could not be obtained due to severe wing deformation, were extracted from the fully-connected (fc)-1000 layer of a pretrained ResNet-50 network (He *et al*. 2015). Fc-1000 is a classification layer that was trained to solve a 1000-way classification problem (He *et al*. 2015) such that the network extracts 1000 features from each image that can be used to train a subsequent machine learning classifier. The pretrained fc-1000 layer was then used to extract wing features from adult wings mounted on glass coverslips. The Classification Learner toolbox in MATLAB was used to train a support-vector machine (Wang 2005) on the labeled data. Fifty images from each of the five representative classes (Crumpled, Doming, Melanotic, Normal, or Thickening) were used for the purpose of training a phenotypic classifier. The approach resulted in a classification accuracy of 97.5% (SI Figure 1).

### *Drosophila* wing quantification

Following the initial separation of wings with severe phenotypes from normal wings, wings classified as containing severe phenotypes underwent qualitative morphological analysis (Figure 1C). Normal wings underwent a separate round of feature extraction to quantify morphology. To extract quantitative features, we used our previously reported open-source pipeline for *Drosophila* wing images. A detailed report on the creation, development, and validation of the pipeline is found in the associated paper on the pipeline (Kumar *et al*. 2022).

Briefly, the pipeline labels the *Drosophila* wing intervein regions using a machine learning-based classifier. *Drosophila* wings have distinct morphological features, including seven intervein regions and the longitudinal veins surrounding them (Figure 1B). Therefore, we trained our analysis algorithm to be able to identify individual intervein regions and extract features accordingly. Geometric features of each intervein region, such as area, aspect ratio, perimeter, eccentricity, and circularity are extracted using MATLAB’s Image Processing Toolbox. These features were then used to train a support-vector machine classifier to label individual intervein regions for all images.

Landmark position-based features, such as the length of the proximal-distal or anterior-posterior axes, were calculated using erosion/dilation operations on the labeled intervein regions to measure longitudinal vein end points. Overall, the pipeline output consists of total wing area, individual intervein region areas, total trichome count, individual intervein trichome count, proximal-distal axis length, anterior-posterior axis length, and the length between the third and fourth longitudinal veins. Quantitative measures using this pipeline were reported to be statistically identical to manual measurements (Kumar *et al*. 2022).

### STRING inference and gene ontology enrichment analysis

We used the Search Tool for the Retrieval of Interacting Genes (STRING) database to investigate known protein-protein interactions (Szklarczyk *et al*. 2015, 2011). The minimum required interaction score was set at a value of 0.700 for high confidence in STRING predictions. The minimum required interaction score is a threshold on the confidence score to screen for positive protein-protein interaction hits. Gene ontology enrichment was performed using PANTHER (Mi *et al*. 2017). The entire *Drosophila* genome was used as the reference list.

### Feature dimension reduction and clustering analysis

In our analysis, we used principal component analysis (PCA) (Pearson 1901), a well-established algorithm to reduce the dimensionality of the data without sacrificing data variation. PCA ensures that distances between individual data points is preserved while projecting the data into lower dimensions. PCA was carried out on wing features extracted by MAPPER (Kumar *et al*. 2022) for wings containing normal morphology. Wing features were aggregated by mean for each gene prior to PCA. PCA components were then subjected to the expectation-maximization algorithm for fitting Gaussian mixture models (EM-GMM) (Yang *et al*. 2012). The EM-GMM algorithm enabled clustering of data points in the principal components using a probabilistic k-means clustering approach (Patel and Kushwaha 2020; Jain 2010).

### Statistical analysis

For the reverse transcription-quantitative polymerase chain reaction experiments, the data was analyzed using R (R Core Team 2022). Associating error bars of the (Δ*R_n_*) plots are representative of the standard error of the mean of each gene at a given cycle count. Experimental runs in which no amplification was detected were given *C_T_* values of 40 as that was the extent of the experimental runs. Pairwise Mann-Whitney U tests were performed for statistical analysis with a false discovery rate (FDR) correction of 0.05 (Benjamini and Hochberg 1995). Reported significance levels are with respect to corrected p-values with raw p-values available in Supplementary File S3.

For the gene ontology enrichment analysis, Fisher’s Exact test (Fisher 1925) was used to identify significant associations of gene ontology. The reported p-value is the corrected values using a false discovery rate of 0.05 to account for potential false positives (Benjamini and Hochberg 1995).

## Results and discussion

### A comprehensive phenotypic map of GPCRs and G proteins in the *Drosophila* wing

All 124 genes were evaluated in their extent to induce either quantitative or qualitative defects in the *Drosophila melanogaster* wing. Of the GPCRs and G proteins screened, 25 demonstrated at least 60% penetrance of severe qualitative phenotypes when knocked down. Observed phenotypes range from mild (incomplete wing veins, bifurcation of the wing veins, and changes in total wing area) to severe (vein thickening, crumpled wings, and melanotic wings) (Figures 2, 3 and SI Figure 2). Candidate genes of interest from the performed screen are defined as MS1096>UAS-RNAi line progeny in which there was at least 40% vein disruption penetrance or at least 60% penetrance of severe phenotypes (see Materials and methods section for details). Using this approach, we identified 29 positive hits that contribute to wing development, with several of the identified genes having low abundance expression reported in the developing *Drosophila* wing disc (Sobala and Adler 2016; Ren *et al*. 2005; Celniker *et al*. 2009). This suggests that low abundance GPCRs may have significant regulatory roles during the morphogenetic process. Of the identified positive hits, 11 genes with some of the most severe phenotypes have viable human orthologs that contribute to a variety of diseases (Table 1). The majority of positive hits identified were from Class A and Class B GPCRs due to the class constituents making up 59% and 20% of the total genes screened, respectively (see Supplementary File S2).

**Figure 2.**
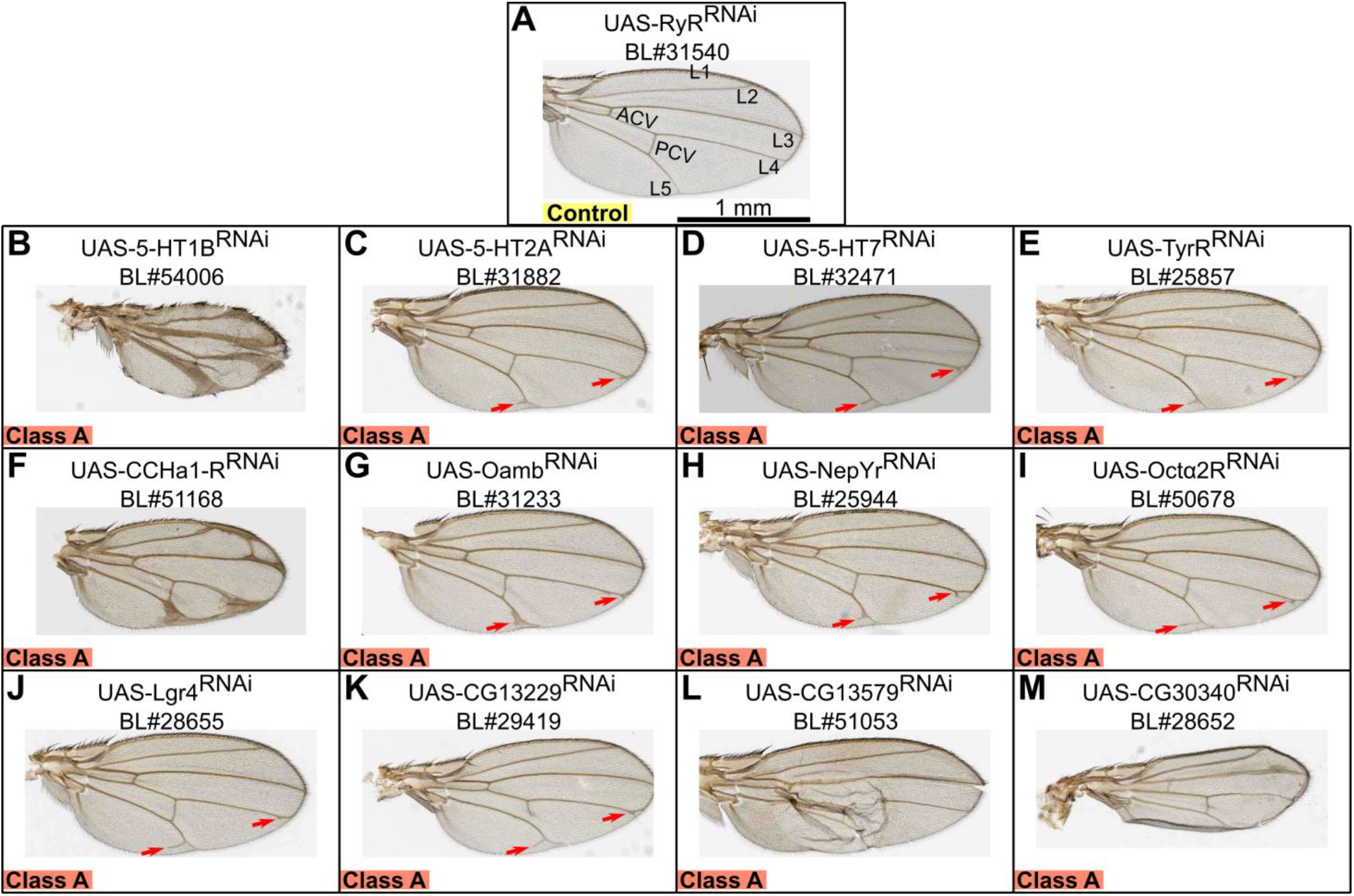
Knockdown of various neuropeptide and neurotransmitter GPCRs results in severe wing phenotypes. Genetic knockdown progeny were generated by crossing the MS1096-Gal4 line to RNAi-based transgenic lines (UAS-Gene *X^RNAi^* (Perkins *et al*. 2015)). The RNAi for the ryanodine receptor (RyR) (BL#31540) was used as the background control for crosses as we have previously demonstrated the MS1096-Gal4 × UAS-RyR^*RNAi*^ cross does not exhibit phenotypic defects due to the RyR gene not being expressed in the *Drosophila* wing disc (Brodskiy *et al*. 2019; Gramates *et al*. 2017). The control group wing **(A)** has five longitudinal veins (L1-L5), an anterior crossvein (ACV), and a posterior crossvein (PCV) without any notable defects. Knockdown of 12 rhodopsin-like (Class A) GPCRs in the developing *Drosophila* wing disc resulted in wing phenotypes ranging from vein bifurcations (red arrows), thickened veins, doming, blistering, and crumpled wings **(B-M)**. The 12 identified GPCRs are either neurotransmitter or neuropeptide receptors whose genes are reported to have low-to-no expression in *Drosophila* wing imaginal discs during larval and pupal stages (Ren *et al*. 2005; Sobala and Adler 2016; Celniker *et al*. 2009). The scale bar in panel **A** represents 1 mm and applies to all panels in the figure. BL# is indicative of the Bloomington *Drosophila* Stock Center stock number for the genetic line used. Images are of adult male wings from F1 progeny resulting from MS1096-Gal4>UAS-Gene *X^RNAi^* crosses.

**Figure 3.**
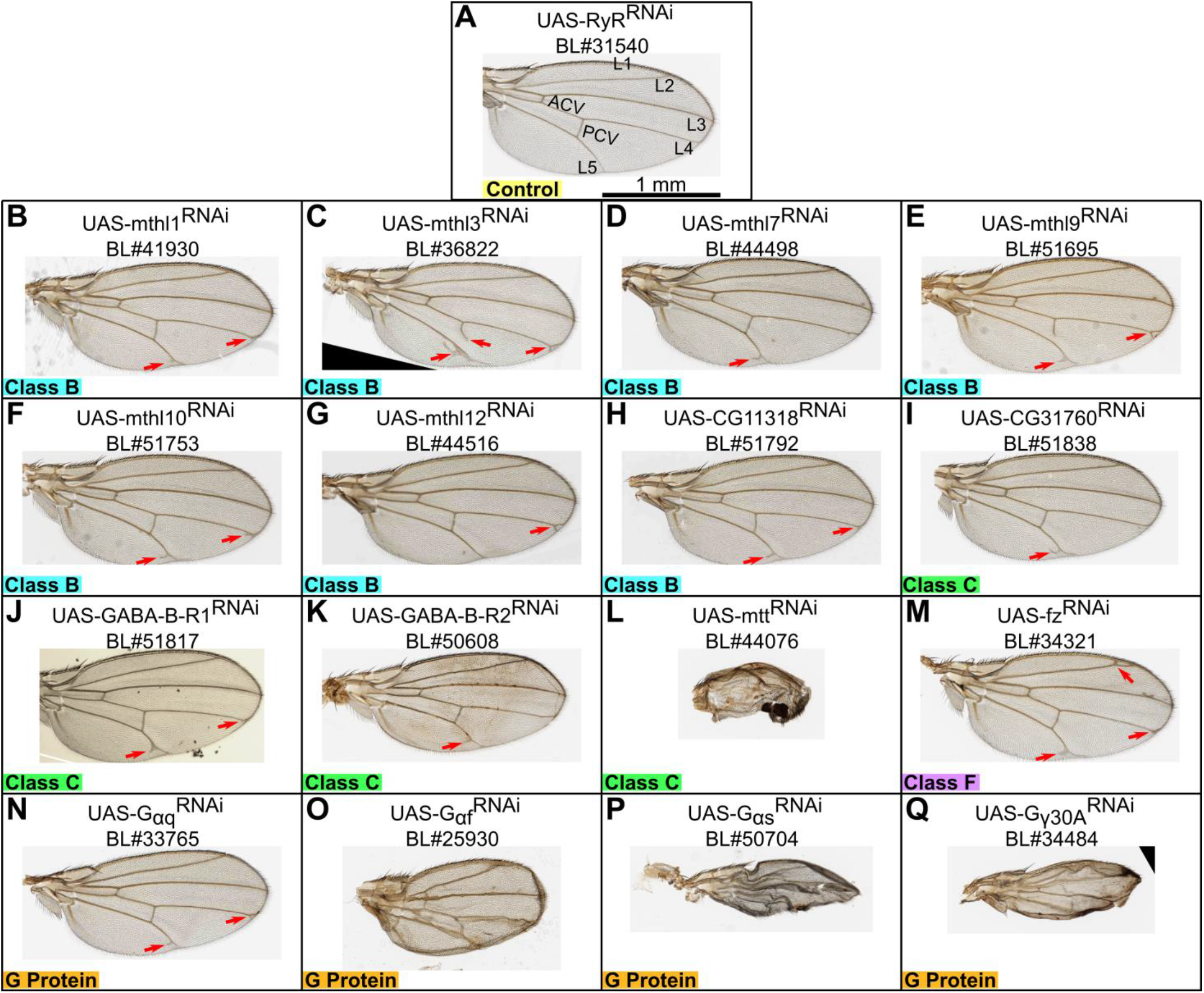
Knockdown of various GPCRs and G proteins results in severe wing phenotypes. The control group wing **(A)** has five longitudinal veins (L1-L5), an anterior crossvein (ACV), and a posterior crossvein (PCV) without notable defects. Knockdown of 7 Class B **(B-H)**, 4 Class C **(I-L)**, 1 Class F **(M)**, and 4 G proteins **(N-Q)** in the developing *Drosophila* wing disc resulted in wing phenotypes ranging from vein bifurcations (red arrows), doming, and melanotic wings. Methuselah (mthl) (Class B) GPCRs consistently produce vein bifurcation defects **(A-F)** while Class C and G protein knockdown has a variety of wing phenotypes. Several of the identified genes **(D,G-L,O,Q)** are reported to have low-to-no expression in *Drosophila* wing imaginal discs during larval and pupal stages (Ren *et al*. 2005; Sobala and Adler 2016; Celniker *et al*. 2009). The scale bar in panel **A** represents 1 mm and applies to all panels in the figure. BL# is indicative of the Bloomington *Drosophila* Stock Center stock number for the genetic line used. Images are of adult male wings from F1 progeny resulting from MS1096-Gal4>UAS-Gene *X^RNAi^* crosses.

Interestingly, knockdown of 12 rhodopsin-like (Class A) GPCRs in the developing *Drosophila* wing disc resulted in wing phenotypes ranging from vein bifurcations, thickened veins, doming, blistering, to crumpled wings (Figure 2). The 12 Class A GPCRs identified are either neurotransmitter or neuropeptide receptors whose genes are reported to have low-to-no expression in the developing *Drosophila* wing imaginal discs during the larval and pupal stages (Table 3) (Sobala and Adler 2016; Ren *et al*. 2005; Celniker *et al*. 2009). Further analysis revealed that knockdown of 7 Class B, 4 Class C, 1 Class F, and 4 G proteins resulted in wing phenotypes ranging from vein bifurcations, doming, and melanotic wings (Figure 3). The Class B Methuselah (mthl) GPCRs consistently produced vein bifurcation defects. Furthermore, several of the identified phenotypes from non-Class A GPCRs resulted from knockdown of genes that are reported to have low-to-no expression in the developing *Drosophila* wing imaginal disc during larval and pupal stages (Table 3) (Sobala and Adler 2016; Ren *et al*. 2005; Celniker *et al*. 2009). These findings suggest that multiple low abundance GPCRs and G proteins lead to severe phenotypic outcomes when dysregulated.

**Table 3.** Various levels of gene expression for neurotransmitter and neuropeptide GPCRs and G proteins in *Drosophila* wings.

| Gene Name | FBgn | GPCR Class | Sobala et al. Expression<sup>a</sup> | Ren et al. Expression<sup>b</sup> | modENCODE Expression<sup>c</sup> | RT-qPCR Expression<sup>d</sup> |
| --- | --- | --- | --- | --- | --- | --- |
| *5-HT1B* | FBgn0263116 | Class A | Moderately high expression | Not reported | No expression | Low expression |
| *5-HT2A* | FBgn0087012 | Class A | High expression | Not reported | No expression | Moderate expression |
| *5-HT7* | FBgn0004573 | Class A | Moderate expression | Not reported | Very low expression | Not Tested |
| *TyrR* | FBgn0038542 | Class A | Moderate expression | Not reported | Very low expression | Not Tested |
| *CCHa1-R* | FBgn0050106 | Class A | Moderately high expression | Not reported | No expression | Low expression |
| *Oamb* | FBgn0024944 | Class A | High expression | Not reported | No expression | Not Tested |
| *NepYr* | FBgn0004842 | Class A | Moderately high expression | Moderate expression | No expression | Not Tested |
| *Octa2R* | FBgn0038653 | Class A | Moderate expression | Not reported | No expression | Not Tested |
| *Lgr4* | FBgn0085440 | Class A | Moderately high expression | Not reported | No expression | Not Tested |
| *CG13229* | FBgn0033579 | Class A | High expression | Not reported | Very low expression | Not Tested |
| *CG13579* | FBgn0035010 | Class A | Moderate expression | Not reported | No expression | Low expression |
| *CG30340* | FBgn0050340 | Class A | Moderate expression | Not reported | No expression | Low expression |
| *mthl1* | FBgn0030766 | Class B | High expression | Not reported | Low expression | Not Tested |
| *mthl3* | FBgn0028956 | Class B | Moderate expression | Not reported | Moderate expression | Moderate expression |
| *mthl7* | FBgn0035847 | Class B | Moderate expression | Not reported | No expression | Not Tested |
| *mthl9* | FBgn0035131 | Class B | High expression | Not reported | Low expression | Not Tested |
| *mthl10* | FBgn0035132 | Class B | High expression | Not reported | Moderate expression | Not Tested |
| *mthl12* | FBgn0045442 | Class B | No expression | Not reported | No expression | Not Tested |
| *CG11318* | FBgn0039818 | Class B | Moderate expression | Not reported | Very low expression | Not Tested |
| *CG31760* | FBgn0051760 | Class C | Moderate expression | Not reported | Very low expression | Not Tested |
| *GABA-B-R1* | FBgn0260446 | Class C | Moderate expression | Not reported | Very low expression | Moderate expression |
| *GABA-B-R2* | FBgn0027575 | Class C | Moderate expression | Not reported | Low expression | Moderate expression |
| *mtt* | FBgn0050361 | Class C | Moderate expression | Not reported | No expression | Very low expression |
| *fz* | FBgn0001085 | Class F | High expression | Not reported | Moderate expression | Not Tested |
| *Gaq* | FBgn0004435 | G protein | High expression | Not reported | Moderately high expression | Not Tested |
| *Gaf* | FBgn0010223 | G protein | Moderate expression | Not reported | Very low expression | Moderate expression |
| *Gas* | FBgn0001123 | G protein | High expression | Not reported | Moderately high expression | Moderate expression |
| *Gγ30A* | FBgn0267252 | G protein | Moderate expression | Not reported | Low expression | Very low expression |

<sup>a</sup> Pupal wing RNAseq from (Sobala and Adler 2016)<sup>b</sup> Pupal wing RNA Affymetrix gene chip from (Ren et al. 2005)<sup>c</sup> Third instar imaginal wing disc RNASeq from (Celniker et al. 2009)<sup>d</sup> Expression from data used to generate Figure 4

### RT-qPCR confirms expression of identified positive hits in developing wing imaginal discs

Select positive hit genes that exhibited severe wing phenotypes were tested to confirm expression in the *Drosophila* imaginal wing disc (Figure 4). To confirm the presence of these GPCRs in the *Drosophila* wing disc, RT-qPCR (Bustin 2000) on third instar *Drosophila* wing disc derived Cl.8 cells (Peel and Milner 1990; Cherbas *et al*. 2011) was performed. Genes for RT-qPCR were selected due to their high penetrance of qualitative phenotypes or due to not having known roles in epithelial morphogenesis. *α*-Tubulin at 84B (NCBI Reference Sequence: NM_057424), a ubiquitously expressed gene, was used as a positive control. No template control (NTC) wells containing no template DNA were used as the negative control. Gene targets of the experiments, their corresponding figure labels, their FlyBase ID, and their NCBI Reference Sequence ID are found in Table 2.

Through observation of the measures Δ*R_n_* and Δ*C_T_* for positive hits, there was a clear indication of mRNA expression, although in low abundance (Figure 4). Of the tested gene targets, G*γ*30A and mtt demonstrated very low, but detectable expression (*p* < 0.01 compared to the negative control). 5-HT1B and CCHa1-R, a neurotransmitter and neuropeptide receptor, respectively, demonstrated low, detectable expression for multiple oligonucleotide primer pairs (*p* < 0.01 compared to the negative control). CG13579 and CG30340, an orphan amine GPCR and neuropeptide receptor, respectively, also demonstrated low, detectable expression (*p* < 0.01 compared to the negative control). The remaining positive hits tested by RT-qPCR demonstrated moderate levels of expression (Figure 4 and Table 3). These data provide new evidence for the expression of low abundance GPCRs that lead to severe phenotypic outcomes when dysregulated. Interestingly, several of these genes were not detected in third instar *Drosophila* wing imaginal discs using alternative RNA-Sequencing methods (Celniker *et al*. 2009). However, others have reported detectable levels of expression in pupal *Drosophila* wings (Sobala and Adler 2016; Ren *et al*. 2005) (Table 3). Literature has reported that the MS1096-Gal4 driver can drive expression in the pupal wing of *Drosophila melanogaster* (Egoz-Matia *et al*. 2011). Therefore, the observed phenotypic defects produced in the RNAi screen may be attributable to either larval or pupal expression of RNAi constructs for the target gene under control of the MS1096-Gal4 driver.

### Quantitative and qualitative analyses highlight the extent of wing phenotypes upon GPCR knockdown

Using our previously reported open-source pipeline for high-content screening of *Drosophila* wing images (MAPPER) (Kumar *et al*. 2022), we looked into how knockdown of GPCRs and G proteins influenced the size of the adult *Drosophila* wing (Figure 5). Due to the severity of the phenotypes produced by knockdown of wings, only wings with standard morphometry were analyzed quantitatively using this pipeline (Figure 1C). Of the 18 GPCRs and G proteins that induced the greatest percent change in wing area compared to the control group (Figure 5A), 12 of the hits were either neurotransmitter or neuropeptide receptors.

**Figure 5.**
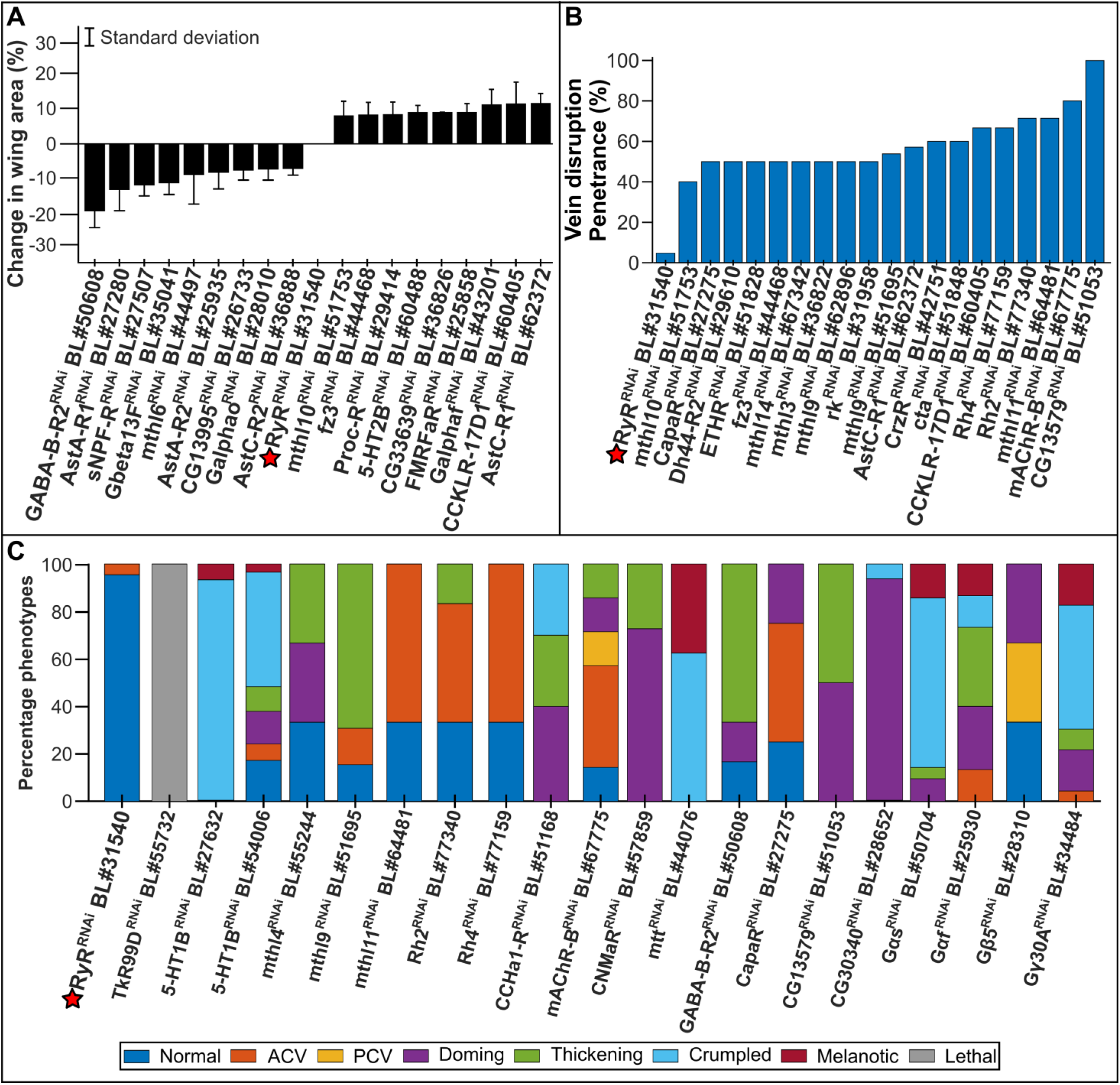
Quantitative and qualitative analyses highlight the extent of wing phenotypes upon GPCR knockdown. **(A)** Change in wing area compared to the control MS1096-Gal4>UAS-RyR^*RNAi*^ is plotted. Only the 18 largest percentage changes in wing area resulting from genetic knockdown are plotted. The error bars represent one standard deviation. **(B)** The penetrance of wings with vein disruption is plotted. Only the top 19 largest percentages of penetration in wing vein disruption resulting from genetic knockdown are plotted. **(C)** Proportions of qualitative phenotypes for genetic knockdown crosses are plotted. ACV: anterior crossvein, PCV: posterior crossvein. Labels are of adult male wings from F1 progeny resulting from MS1096-Gal4>UAS-Gene *X^RNAi^* crosses.

For wings with less severe qualitative phenotypes, MAPPER was able to assist in quantifying the penetrance of vein disruption (Figure 5B). Vein disruption was quantified as wings containing crossvein defects (SI Figure 2C,D), bifurcation defects (Figure 3B-K), or blistering defects (Figure 2L). The morphology of the veins is precisely regulated by multiple morphogenetic pathways, including Decapentaplegic (Dpp), Hedgehog, EGFR, and Notch (Blair 2007; Matsuda and Shimmi 2012; Ralston and Blair 2005). Therefore, the presence of vein disruption may provide insight as to how GPCRs and G proteins interact with morphogenetic signaling pathways. Of the 19 GPCRs and G proteins that induced the greatest percent change in vein disruption compared to the control group (Figure 5B), 10 of the hits were Class A GPCRs. Furthermore, several of the greatest penetrance outcomes of vein disruption were the result of the knockdown of Methuselah receptors. Of the 19 GPCRs and G proteins that induced the greatest percent change in vein disruption, 6 of the hits were members of the Methuselah family. Interestingly, there is evidence demonstrating the role of Methuselah and its ligand in regulating epithelial morphogenesis (Manning *et al*. 2013), thereby giving credence to the role of GPCRs in morphogenesis.

For wings with the greatest phenotypic defects, qualitative features of the wing data was extracted from the fully-connected (fc)-1000 layer of a pretrained ResNet-50 network (He *et al*. 2015). A support-vector machine (Wang 2005) was then trained to classify the qualitative features into five representative classes: Crumpled, Doming, Melanotic, Normal, or Thickening (SI Figure 2). The trained support-vector machine resulted in a classification accuracy of 97.5% (SI Figure 1). We observed a large range of penetrance in defects among the identified genes wherein several GPCR knockdowns, such as TkR99D (BL#55732), CCHa1-R (BL#51168), and 5-HT1B (BL#27632), resulted in 100% penetrance of severe wing phenotypes while other knockdowns resulted in moderate venation defects (Figure 5C).

### Gaussian mixture models and Euclidean distances unveil predictions for unreported protein-protein interactions

To evaluate the quantitative phenotypes more comprehensively, morphological wing features extracted from MAPPER (Kumar *et al*. 2022) were mapped to a two-dimensional space using principal component analysis. Four clusters were identified using Gaussian mixture models and Bayesian information criterion (BIC) with each cluster containing distinct traits (Figure 6A and SI Figure 3). Constituents of Cluster 1 had larger wings on average (1.084 mm^2^) compared to the wings of Cluster 2 (0.999 mm^2^, *p* = 8.94×10^−16^ via two-tail paired t-test). Further, constituents from Cluster 3 included wings with anterior crossvein defects while Cluster 4 consisted of wings with posterior crossvein defects.

**Figure 6.**
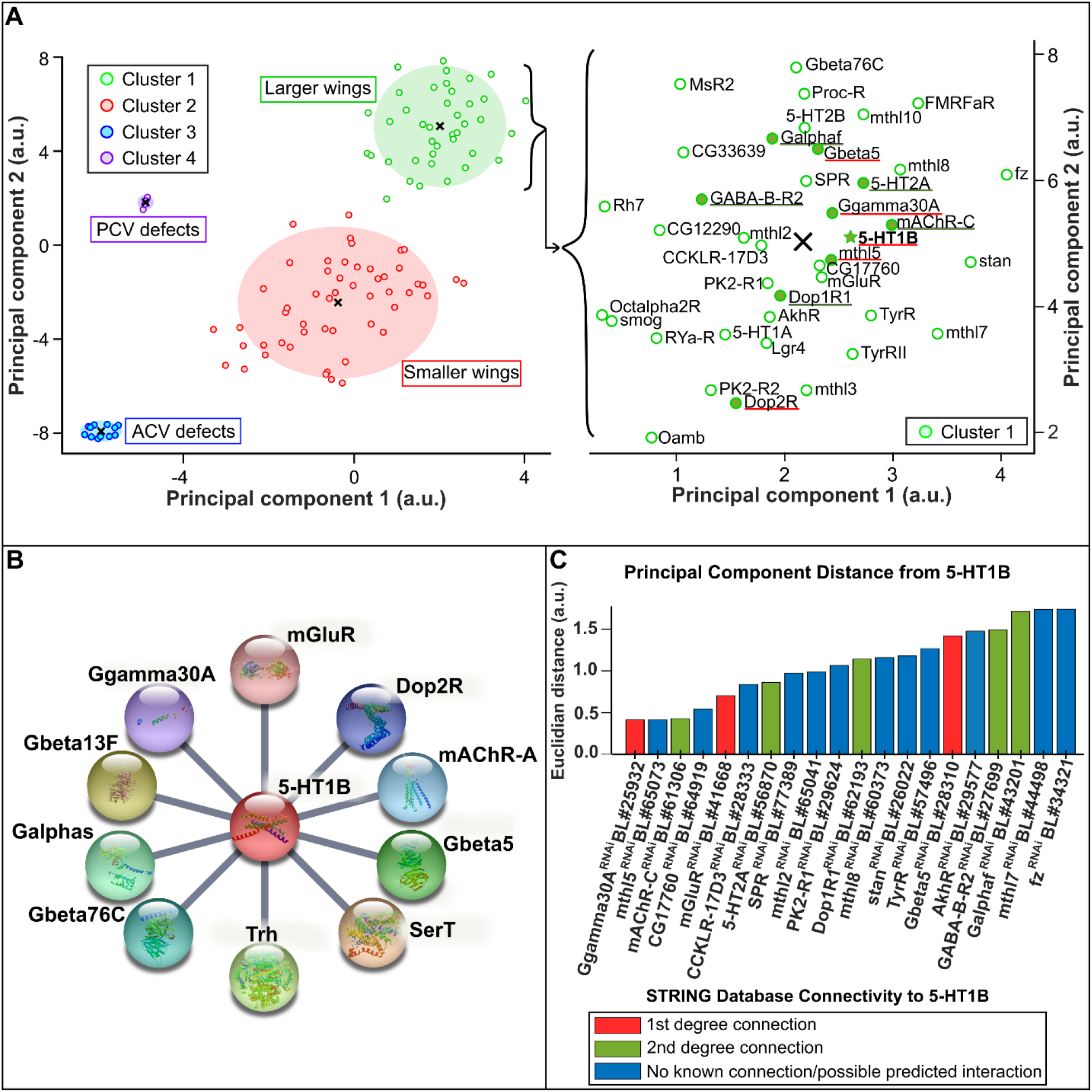
Gaussian mixture models and Euclidean distances unveil predictions for unreported protein-protein interactions. **(A)** Morphological wing features extracted from MAPPER (Kumar *et al*. 2022) were mapped to a two-dimensional space using principal component analysis for analyzed wings resulting from genetic knockdown experiments. Four clusters were identified using Gaussian mixture models with each cluster containing distinct traits. The constituents of Cluster 1 (right) consist of neurotransmitter and neuropeptide receptors, with the 5-HT1B receptor highlighted (green star). Centers of the cluster (x-marks) are representative of the mean properties of the particular cluster. ACV: anterior crossvein, PCV: posterior crossvein. **(B)** The STRING protein-protein interaction network of 5-HT1B is shown (Szklarczyk *et al*. 2015) to depict known protein-protein interactions. **(C)** Protein-protein interactions are predicted using Euclidean distances. The Euclidean distance between 5-HT1B and other genetic knockdowns in the principal component space is plotted for the 20 smallest Euclidean distances (eight known from STRING and 12 novel predictions). Highlighted protein-protein interaction connections are underlined in panel **A**, with the color corresponding to the degree of connection (either 1st in red or 2nd in green).

Because neurotransmitter and neuropeptide receptors consistently produced severe wing phenotypes and were detectable via RT-qPCR experiments, we performed a gene ontology enrichment analysis for the top 20 genes screened that resulted in the greatest penetrance of phenotypes. The resulting gene ontology term of this analysis was “neuropeptide signaling pathway” with a 70.24-fold enrichment (*p* = 9.17×10 ^6^) (Ashburner *et al*. 2000; The Gene Ontology Consortium 2019). With this in mind, we further looked at the constituents of Cluster 1 that consisted mainly of neurotransmitter and neuropeptide receptors (Figure 6A). Near the center of the cluster is the neurotransmitter receptor, 5-HT1B, wherein the center of each cluster is representative of the mean properties of the cluster.

5-HT1B is a well-studied neural receptor that relays information passed on by neurotransmitters (Nebigil *et al*. 2001). Several studies have looked at serotonin as a regulator of early embryogenesis and morphogenesis (Lauder *et al*. 1988; Choi *et al*. 1997; Buznikov *et al*. 2001). However, these studies did not fully explore the role of serotonin in epithelial tissue development. To further understand the potential role of 5-HT1B in morphogenesis, we utilized the STRING protein-protein interaction network (Szklarczyk *et al*. 2011, 2015) to map out known protein-protein interactions of 5-HT1B (Figure 6B). The STRING protein-protein interaction network demonstrates that 5-HT1B most closely interacts with G proteins and other GPCRs.

Because several of the interactions in the STRING network with 5-HT1B consisted of genes that were screened in this study, we measured the Euclidean distance between 5-HT1B and the other genes in the reduced dimensional space (Figure 6A,C). The lower the value of the Euclidean distance, the closer the two genes are in the principal component space, and the higher likelihood of protein-protein interaction as the genes produce morphometirc wing phenotypes of high similarity. Therefore, the Euclidean distance was used as a predictor of protein-protein interactions. Of the 20 lowest Euclidean distances measured, 8 correspond to known protein-protein interactions (either 1st- or 2nd-degree connections) in the STRING database. These results suggest that Euclidean distance in a reduced dimensional space may serve as a simplified way to uncover potential protein-protein interactions through phenotypic analysis.

## Discussion

Here, we have reported a systematic RNAi-based investigation into phenotypes associated with knockdown of various GPCRs and G proteins during *Drosophila melanogaster* wing development. A coupled machine learning-based approach was used to both quantitatively and qualitatively analyze phenotypes. Utilizing the double layered analyses, we discovered several GPCRs and G proteins that demonstrate severe phenotypes when knocked down. Of the 111 GPCRs and 13 G proteins screened with RNAi, we identified 29 positive hits that contribute to wing development. Positive hits contain at least 60%penetrance of severe qualitative phenotypes or at least 40% penetrance of vein disruption.

Although we identified 29 positive hits in the RNAi screen, the reported results very likely underestimate the true impact of GPCRs and G proteins involved in *Drosophila* wing development. From the qualitative analysis, we observed a wide range of penetrance in defects among positive hits. This variability in penetrance may be attributable to several of the screened GPCRs demonstrating low abundance expression in the developing *Drosophila* wing disc (Table 3) (Sobala and Adler 2016; Ren *et al*. 2005; Celniker *et al*. 2009). Therefore, severe phenotypes may be observed due to knocking down the already minuscule levels of expression. An additional explanation in the variability of penetrance may be due to various gene paralogs playing compensatory roles when there is reduced function of a specific GPCR. This effect has been observed in ion channel depletion wherein Irk2 disruption is compensated for by increased Irk1 and Irk3 expression (Dahal *et al*. 2012; George *et al*. 2019). Due to the functional crosstalk that occurs between GPCRs (Hur and Kim 2002; Guo *et al*. 2005; Saad and Hipfner 2021), GPCR compensatory mechanisms may rescue more severe defects from occurring.

Several of the identified hits were neurotransmitter or neuropeptide receptors not known to play roles in epithelial morphogenesis. However, because there is conflicting evidence for either no expression or weak expression patterns in the central nervous system when using the MS1096-Gal4 driver (Lindström *et al*. 2017; Ray and Lakhotia 2019), identified hits require further characterization of knockdown phenotypes under the control of other Gal4 lines. If the utilization of other Gal4 lines confirms similar reported phenotypes, positive hits would warrant further investigation to validate the genes as potential disease models. However, if there are no phenotypes or discrepant phenotypes observed under the control of other Gal4 drivers, the results presented here may be attributable to cell non-autonomous roles of GPCRs and G proteins in the nervous system. Literature reports demonstrate the crucial role of the developing brain and peripheral nerves in coordinating morphogenesis of surrounding tissues (Adameyko and Fried 2016). Due to the extensive crosstalk involved with GPCRs and morphogenetic signaling pathways, such as Hedgehog (Saad and Hipfner 2021), nervous system dysregulation by GPCRs could lead to non-cell autonomous aberrations in other developing tissues.

RT-qPCR experiments confirmed the presence, although in low abundance, of several of the positive hits. Interestingly, many of the identified hits have not been reported to have third instar larval wing disc expression (Celniker *et al*. 2009), despite having detectable levels of expression in pupal wings (Sobala and Adler 2016; Ren *et al*. 2005) (Table 3). The MS1096-Gal4 driver has been used to drive gene expression in pupal wings (Egoz-Matia *et al*. 2011). Therefore, future experimentation is required to determine if the reported phenotypes are primarily due to larval or pupal expression of the RNAi constructs under control of MS1096-Gal4. However, because the RT-qPCR experiments were performed on third instar larval wing disc cells, we deduce that phenotypic outcomes are likely a result in part of larval expression of the RNAi constructs.

Further analysis of neurotransmitter and neuropeptide GPCR hits demonstrated that Euclidean distances of quantitative wing features can be used to predict protein-protein interactions. By measuring the Euclidean distance between the neurotransmitter receptor, 5-HT1B, and other genes, we predicted 20 potential protein-protein interactions (Figure 6). Eight of the predicted interactions were confirmed as 1st- or 2nd-degree connections using the STRING database (Szklarczyk *et al*. 2011, 2015). The remaining 12 distances not validated by STRING may be inferred as novel predictions for unreported protein-protein interactions with 5-HT1B that require further elucidation. For example, a recent study (Malpe *et al*. 2020) demonstrated that *5-HT1B* and *mthl-5* expression are both required for mitosis in germline stem cell division in *Drosophila*. This discovery provides additional evidence for the Euclidean distance predicted protein-protein interaction between 5-HT1B and mthl-5 (Figure 6C).

Although many of the identified positive hits are known for their roles in regulating neural activity, very little is known about the role of such GPCRs in *Drosophila* development (Hanlon and Andrew 2015; Manning *et al*. 2013; Schwabe *et al*. 2005; Hannon and Hoyer 2008; Terunuma 2018; Barnes and Sharp 1999). To the best of our knowledge, our study is the first to utilize a coupled quantitative and qualitative machine learning-based approach to systematically investigate the role of GPCRs in *Drosophila* wing development. Therefore, the role of the identified GPCRs and G proteins in epithelial morphogenesis warrants further investigation. Future studies can lead to the development of in vivo models for diseases associated with dysregulation of the *Drosophila* gene human orthologs (Table 1).

Interestingly, the phenotypic manifestations in humans for identified orthologs have similar phenotypes reported from the RNAi screen. For example, knockdown of *5-HT1B, Rh2, CCHa1-R*, and *fz* in the developing *Drosophila* wing produced vein thickening phenotypes. The human orthologs for these genes, *HTR1A, OPN4, GRPR*, and *FZD7*, respectively, have disease manifestations that increase metastasis, tumor cell growth, cell migration, and endothelial cell proliferation (Table 1) (Liu *et al*. 2022a; de Assis *et al*. 2021; Patel *et al*. 2014; Zhang and Xu 2022). Similar vein thickening defects have been observed when knocking down the *dSTIM* gene, a calcium release-activated calcium (CRAC) channel (Eid *et al*. 2008). Literature reports have demonstrated the role of enhanced mobilization of intracellular calcium resulting from crosstalk between different GPCRs (Werry *et al*. 2003). Therefore, vein thickening defects observed after knockdown of GPCRs may be the result of disrupted intracellular calcium regulation.

Overall, the combined machine learning approaches for both qualitative and quantitative analyses enabled a more comprehensive analysis of the 29 identified GPCRs and G proteins that may regulate *Drosophila* wing development. The results of this study provide a starting point for further exploration of the signaling pathways and molecular mechanisms underlying GPCR regulation and cross-talk during epithelial morphogenesis.

## Data availability

A comprehensive list of genes screened and associated Bloomington *Drosophila* Stock Center numbers is provided in File S1. A comprehensive list of the screened genes, their FlyBase IDs, their GPCR class, and general gene information is provided in Supplementary File S2. Supplementary File S3 provides the raw and corrected p-values for the RT-qPCR experiments, while Supplementary File S4 provides the raw data generated by the StepOne^*TM*^ software for the RT-qPCR experiments. Supplementary File S5 provides wing data information used to generate Figures 5 and 6.

## Acknowledgments

We would like to thank the South Bend Medical Foundation for granting generous access to their Apero Slide Scanner, and Kara Snyder, Mary Kate O’Leary, Seth Tautges, Heather Flynn, Ryan Govi, Quincey Hogue,

Lev S. Suliandziga, Spencer Hayes, Adriana Szpynda, Isaac Nordan, Mark Legendre, Dharsan Soundarrajan, Czier-Anne Gone, Meaghan Snyder, Andrew Blake, Ulises Hernandez Martin DelCampo, Brooke Gleason, Cesar Moreno, Eleanora Korotkova, and Svitlana Bychko for technical assistance in performing the screen. Additionally, we thank Antonio Loreto and Daniela D. Ruedas for their assistance in the curation of human orthologs of positive hit genes from the *Drosophila* screen. We thank Mayesha S. Mim and David Gazzo for valuable feedback and discussion of the manuscript.

## Literature cited

Adameyko I, Fried K. 2016. The Nervous System Orchestrates and Integrates Craniofacial Development: A Review. Frontiers in Physiology. 7:49.

Adams JC, Watt FM. 1993. Regulation of development and differentiation by the extracellular matrix. Development (Cambridge, England). 117:1183–1198.

Adams JU. 2014. G-Protein-Coupled Receptors Play Many Different Roles in Eukaryotic Cell Signaling, In: O’Connor C, editor, Essentials of Cell Biology, Nature Education. How do cells sense their environment?. Cg_cat: GPCR Cg_level: ESY Cg_topic: GPCR.

Ashburner M, Ball CA, Blake JA, Botstein D, Butler H, Cherry JM, Davis AP, Dolinski K, Dwight SS, Eppig JT et al. 2000. Gene Ontology: tool for the unification of biology. Nature Genetics. 25:25–29.

Ashburner M, Golic K, Hawley R. 2005. Drosophila: A Laboratory Handbook. Cold Spring Harbor Laboratory Press. Cold Spring Harbour, NY.

Ayers KL, Thérond PP. 2010. Evaluating Smoothened as a G-Protein-Coupled Receptor for Hedgehog Signalling. Trends in Cell Biology. 20:287–298.

Barnes NM, Sharp T. 1999. A review of central 5-HT receptors and their function. Neuropharmacology. 38:1083–1152.

Bastiaansen AEM, Timmermans AM, Smid M, van Deurzen CHM, Hulsenboom ESP, Prager-van der Smissen WJC, Foekens R, Trapman-Jansen AMAC, Sillevis Smitt PAE, Luider TM et al. 2020. Metabotropic glutamate receptor 1 is associated with unfavorable prognosis in ER-negative and triple-negative breast cancer. Scientific Reports. 10:22292.

Belacortu Y, Paricio N. 2011. Drosophila as a model of wound healing and tissue regeneration in vertebrates. Developmental Dynamics. 240:2379–2404.

Belgacem YH, Borodinsky LN. 2011. Sonic hedgehog signaling is decoded by calcium spike activity in the developing spinal cord. Proceedings of the National Academy of Sciences. 108:4482–4487.

Benjamini Y, Hochberg Y. 1995. Controlling the False Discovery Rate: A Practical and Powerful Approach to Multiple Testing. Journal of the Royal Statistical Society. Series B (Methodological). 57:289–300. Publisher: [Royal Statistical Society, Wiley].

Bettler B, Kaupmann K, Mosbacher J, Gassmann M. 2004. Molecular structure and physiological functions of GABA(B) receptors. Physiological Reviews. 84:835–867.

Blair SS. 2007. Wing Vein Patterning in Drosophila and the Analysis of Intercellular Signaling. Annual Review of Cell and Developmental Biology. 23:293–319.

Blau HM, Baltimore D. 1991. Differentiation requires continuous regulation. The Journal of Cell Biology. 112:781–783.

Brodskiy PA, Wu Q, Soundarrajan DK, Huizar FJ, Chen J, Liang P, Narciso C, Levis MK, Arredondo-Walsh N, Chen DZ et al. 2019. Decoding Calcium Signaling Dynamics during Drosophila Wing Disc Development. Biophysical Journal. 116:725–740.

Buchmann A, Alber M, Zartman JJ. 2014. Sizing it up: The mechanical feedback hypothesis of organ growth regulation. Seminars in Cell & Developmental Biology. 35:73–81.

Bustin SA. 2000. Absolute quantification of mRNA using real-time reverse transcription polymerase chain reaction assays. Journal of Molecular Endocrinology. 25:169–193.

Buznikov GA, Lambert HW, Lauder JM. 2001. Serotonin and serotonin-like substances as regulators of early embryogenesis and morphogenesis. Cell and Tissue Research. 305:177–186.

Capdevila J, Guerrero I. 1994. Targeted expression of the signaling molecule decapentaplegic induces pattern duplications and growth alterations in Drosophila wings. The EMBO journal. 13:4459–4468.

Carecchio M, Panteghini C, Reale C, Barzaghi C, Monti V, Romito L, Sasanelli F, Garavaglia B. 2016. Novel GNAL mutation with intra-familial clinical heterogeneity: Expanding the phenotype. Parkinsonism & Related Disorders. 23:66–71.

Celniker SE, Dillon LAL, Gerstein MB, Gunsalus KC, Henikoff S, Karpen GH, Kellis M, Lai EC, Lieb JD, MacAlpine DM et al. 2009. Unlocking the secrets of the genome. Nature. 459:927–930.

Cherbas L. 2016. Additions to Tissue Culture Medium.

Cherbas L, Willingham A, Zhang D, Yang L, Zou Y, Eads BD, Carlson JW, Landolin JM, Kapranov P, Dumais J et al. 2011. The transcriptional diversity of 25 Drosophila cell lines. Genome Research. 21:301–314. Publisher: Cold Spring Harbor Lab.

Choi DS, Ward SJ, Messaddeq N, Launay JM, Maroteaux L. 1997. 5-HT2B receptor-mediated serotonin morphogenetic functions in mouse cranial neural crest and myocardiac cells. Development (Cambridge, England). 124:1745–1755.

Dahal GR, Rawson J, Gassaway B, Kwok B, Tong Y, Ptácěk LJ, Bates E. 2012. An inwardly rectifying K+ channel is required for patterning. Development (Cambridge, England). 139:3653–3664.

De Assis LVM, Lacerda JT, Moraes MN, Domínguez-Amorocho OA, Kinker GS, Mendes D, Silva MM, Menck CFM, Câmara NOS, Castrucci AMdL. 2022. Melanopsin (Opn4) is an oncogene in cutaneous melanoma. Communications Biology. 5:461.

De Assis LVM, Moraes MN, Mendes D, Silva MM, Menck CFM, Castrucci AMdL. 2021. Loss of Melanopsin (OPN4) Leads to a Faster Cell Cycle Progression and Growth in Murine Melanocytes. Current Issues in Molecular Biology. 43:1436–1450.

De Nittis P, Efthymiou S, Sarre A, Guex N, Chrast J, Putoux A, Sultan T, Raza Alvi J, Ur Rahman Z, Zafar F et al. 2021. Inhibition of G-protein signalling in cardiac dysfunction of intellectual developmental disorder with cardiac arrhythmia (IDDCA) syndrome. Journal of Medical Genetics. 58:815–831.

Deutschländer AB, Wszolek ZK. 1993. DYT-GNAL, In: Adam MP, Everman DB, Mirzaa GM, Pagon RA, Wallace SE, Bean LJ, Gripp KW, Amemiya A, editors, GeneReviews®, University of Washington, Seattle. Seattle (WA).

Duffy JB. 2002. GAL4 system in Drosophila: a fly geneticist’s Swiss army knife. Genesis (New York, N.Y.: 2000). 34:1–15.

Egoz-Matia N, Nachman A, Halachmi N, Toder M, Klein Y, Salzberg A. 2011. Spatial regulation of cell adhesion in the Drosophila wing is mediated by Delilah, a potent activator of *β*PS integrin expression. Developmental Biology. 351:99–109.

Eid JP, Arias AM, Robertson H, Hime GR, Dziadek M. 2008. The DrosophilaSTIM1 orthologue, dSTIM, has roles in cell fate specification and tissue patterning. BMC Developmental Biology. 8:104.

Etournay R, Popovic’ M, Merkel M, Nandi A, Blasse C, Aigouy B, Brandl H, Myers G, Salbreux G, Jülicher F et al. 2015. Interplay of cell dynamics and epithelial tension during morphogenesis of the Drosophila pupal wing. eLife. 4:e07090. Publisher: eLife Sciences Publications, Ltd.

Fisher SRA. 1925. Statistical Methods for Research Workers. Oliver and Boyd. Google-Books-ID: I0NBAAAAIAAJ.

Foord SM, Bonner TI, Neubig RR, Rosser EM, Pin JP, Davenport AP, Spedding M, Harmar AJ. 2005. International Union of Pharmacology. XLVI. G Protein-Coupled Receptor List. Pharmacological Reviews. 57:279–288. Publisher: American Society for Pharmacology and Experimental Therapeutics Section: Article.

Fristrom D. 1988. The cellular basis of epithelial morphogenesis. A review. Tissue and Cell. 20:645–690.

Garcia-Alloza M, Hirst WD, Chen CPLH, Lasheras B, Francis PT, Ramírez MJ. 2004. Differential Involvement of 5-HT1B/1D and 5-HT6 Receptors in Cognitive and Non-cognitive Symptoms in Alzheimer’s Disease. Neuropsychopharmacology. 29:410–416. Number: 2 Publisher: Nature Publishing Group.

Garcia De Las Bayonas A, Philippe JM, Lellouch AC, Lecuit T. 2019. Distinct RhoGEFs Activate Apical and Junctional Contractility under Control of G Proteins during Epithelial Morphogenesis. Current biology: CB. 29:3370–3385.e7.

Garczyk S, Klotz N, Szczepanski S, Denecke B, Antonopoulos W, von Stillfried S, Knüchel R, Rose M, Dahl E. 2017. Oncogenic features of neuromedin U in breast cancer are associated with NMUR2 expression involving crosstalk with members of the WNT signaling pathway. Oncotarget. 8:36246–36265.

George LF, Pradhan SJ, Mitchell D, Josey M, Casey J, Belus MT, Fedder KN, Dahal GR, Bates EA. 2019. Ion Channel Contributions to Wing Development in Drosophila melanogaster. G3: Genes|Genomes|Genetics. 9:999–1008.

Ghosh E, Kumari P, Jaiman D, Shukla AK. 2015. Methodological advances: the unsung heroes of the GPCR structural revolution. Nature Reviews Molecular Cell Biology. 16:69–81. Number: 2 Publisher: Nature Publishing Group.

Gramates LS, Marygold SJ, dos Santos G, Urbano JM, Antonazzo G, Matthews BB, Rey AJ, Tabone CJ, Crosby MA, Emmert DB et al. 2017. FlyBase at 25: looking to the future. Nucleic Acids Research. 45:D663–D671.

Guo W, Shi L, Filizola M, Weinstein H, Javitch JA. 2005. Crosstalk in G protein-coupled receptors: Changes at the transmembrane homodimer interface determine activation. Proceedings of the National Academy of Sciences. 102:17495–17500. Publisher: Proceedings of the National Academy of Sciences.

Hanlon CD, Andrew DJ. 2015. Outside-in signaling – a brief review of GPCR signaling with a focus on the Drosophila GPCR family. Journal of Cell Science. 128:3533–3542.

Hannon J, Hoyer D. 2008. Molecular biology of 5-HT receptors. Behavioural Brain Research. 195:198–213.

He K, Zhang X, Ren S, Sun J. 2015. Deep Residual Learning for Image Recognition. arXiv:1512.03385 [cs].

Heigwer F, Port F, Boutros M. 2018. RNA Interference (RNAi) Screening in Drosophila. Genetics. 208:853–874.

Hepler JR, Gilman AG. 1992. G proteins. Trends in Biochemical Sciences. 17:383–387.

Hu Y, Flockhart I, Vinayagam A, Bergwitz C, Berger B, Perrimon N, Mohr SE. 2011. An integrative approach to ortholog prediction for disease-focused and other functional studies. BMC Bioinformatics. 12:357.

Huang Y, Skwarek-Maruszewska A, Horré K, Vandewyer E, Wolfs L, Snellinx A, Saito T, Radaelli E, Corthout N, Colombelli J et al. 2015. Loss of GPR3 reduces the amyloid plaque burden and improves memory in Alzheimer’s disease mouse models. Science Translational Medicine. 7:309ra164.

Hur EM, Kim KT. 2002. G protein-coupled receptor signalling and cross-talk: achieving rapidity and specificity. Cellular Signalling. 14:397–405.

Insel PA, Sriram K, Gorr MW, Wiley SZ, Michkov A, Salmerón C, Chinn AM. 2019. GPCRomics: An approach to discover GPCR drug targets. Trends in pharmacological sciences. 40:378–387.

Jain AK. 2010. Data clustering: 50 years beyond K-means. Pattern Recognition Letters. 31:651–666.

Jennings BH. 2011. Drosophila – a versatile model in biology & medicine. Materials Today. 14:190–195.

Khan SM, Sleno R, Gora S, Zylbergold P, Laverdure JP, Labbé JC, Miller GJ, Hébert TE. 2013. The Expanding Roles of *Gβγ* Subunits in G Protein–Coupled Receptor Signaling and Drug Action. Pharmacological Reviews. 65:545–577.

Kopparapu PK, Tinzl M, Anagnostaki L, Persson JL, Dizeyi N. 2013. Expression and localization of serotonin receptors in human breast cancer. Anticancer Research. 33:363–370.

Kumar N, Huizar FJ, Farfán-Pira KJ, Brodskiy PA, Soundarrajan DK, Nahmad M, Zartman JJ. 2022. MAPPER: An Open-Source, High-Dimensional Image Analysis Pipeline Unmasks Differential Regulation of Drosophila Wing Features. Frontiers in Genetics. 13.

Lauder JM, Tamir H, Sadler TW. 1988. Serotonin and morphogenesis. I. Sites of serotonin uptake and -binding protein immunoreactivity in the midgestation mouse embryo. Development (Cambridge, England). 102:709–720.

LeCun Y, Bengio Y, Hinton G. 2015. Deep learning. Nature. 521:436–444. Number: 7553 Publisher: Nature Publishing Group.

Lee Y, Basith S, Choi S. 2018. Recent Advances in Structure-Based Drug Design Targeting Class A G Protein-Coupled Receptors Utilizing Crystal Structures and Computational Simulations. Journal of Medicinal Chemistry. 61:1–46. Publisher: American Chemical Society.

Lin DM, Goodman CS. 1994. Ectopic and increased expression of Fasciclin II alters motoneuron growth cone guidance. Neuron. 13:507–523.

Lin TY, Huang WL, Lee WY, Luo CW. 2015. Identifying a Neuromedin U Receptor 2 Splice Variant and Determining Its Roles in the Regulation of Signaling and Tumorigenesis In Vitro. PloS One. 10:e0136836.

Lindström R, Lindholm P, Palgi M, Saarma M, Heino TI. 2017. In vivo screening reveals interactions between Drosophila Manf and genes involved in the mitochondria and the ubiquinone synthesis pathway. BMC Genetics. 18:52.

Liu Q, Sun H, Liu Y, Li X, Xu B, Li L, Jin W. 2022a. HTR1A Inhibits the Progression of Triple-Negative Breast Cancer via TGF-*α* Canonical and Noncanonical Pathways. Advanced Science. 9:2105672.

Liu Y, Gu S, Wu T, Xu Y, Meng Q, Guan Z, Zhang X, Sun X, Shen Y, Fan H et al. 2022b. High GNG13 expression is associated with poor survival in epithelial ovarian cancer and breast cancer. Neoplasma. 69:183–192.

Luhur A, Klueg KM, Zelhof AC. 2019. Generating and working with Drosophila cell cultures: Current challenges and opportunities. Wiley interdisciplinary reviews. Developmental biology. 8:e339.

López-Figueroa AL, Norton CS, López-Figueroa MO, Armellini-Dodel D, Burke S, Akil H, López JF, Watson SJ. 2004. Serotonin 5-HT1A, 5-HT1B, and 5-HT2A receptor mRNA expression in subjects with major depression, bipolar disorder, and schizophrenia. Biological Psychiatry. 55:225–233.

López-Varea A, Ostalé CM, Vega-Cuesta P, Ruiz-Gómez A, Organista MF, Martín M, Hevia CF, Molnar C, de Celis J, Culi J et al. 2021a. Genome-wide phenotypic RNAi screen in the Drosophila wing: global parameters. G3 Genes|Genomes|Genetics. 11:jkab351.

López-Varea A, Vega-Cuesta P, Ruiz-Gómez A, Ostalé CM, Molnar C, Hevia CF, Martín M, Organista MF, de Celis J, Culí J et al. 2021b. Genome-wide phenotypic RNAi screen in the Drosophila wing: phenotypic description of functional classes. G3 (Bethesda, Md.). 11:jkab349.

Malpe MS, McSwain LF, Kudyba K, Ng CL, Nicholson J, Brady M, Qian Y, Choksi V, Hudson AG, Parrott BB et al. 2020. G-protein signaling is required for increasing germline stem cell division frequency in response to mating in Drosophila males. Scientific Reports. 10:3888.

Manning AJ, Peters KA, Peifer M, Rogers SL. 2013. Regulation of Epithelial Morphogenesis by the G Protein–Coupled Receptor Mist and Its Ligand Fog. Sci. Signal.. 6:ra98–ra98.

Matsuda S, Shimmi O. 2012. Directional transport and active retention of Dpp/BMP create wing vein patterns in Drosophila. Developmental Biology. 366:153–162.

Mi H, Huang X, Muruganujan A, Tang H, Mills C, Kang D, Thomas PD. 2017. PANTHER Version 11: Expanded Annotation Data from Gene Ontology and Reactome Pathways, and Data Analysis Tool Enhancements. Nucleic Acids Research. 45:D183–D189.

Michelli M, Zougros A, Chatziandreou I, Michalopoulos NV, Lazaris AC, Saetta AA. 2020. Concurrent Wnt pathway component expression in breast and colorectal cancer. Pathology, Research and Practice. 216:153005.

Mirth C, Shingleton A. 2012. Integrating Body and Organ Size in Drosophila: Recent Advances and Outstanding Problems. Frontiers in Endocrinology. 3.

Mirzoyan Z, Sollazzo M, Allocca M, Valenza AM, Grifoni D, Bellosta P. 2019. Drosophila melanogaster: A Model Organism to Study Cancer. Frontiers in Genetics. 10.

Morgat C, MacGrogan G, Brouste V, Vélasco V, Sévenet N, Bonnefoi H, Fernandez P, Debled M, Hindié E. 2017. Expression of Gastrin-Releasing Peptide Receptor in Breast Cancer and Its Association with Pathologic, Biologic, and Clinical Parameters: A Study of 1,432 Primary Tumors. Journal of Nuclear Medicine: Official Publication, Society of Nuclear Medicine. 58:1401–1407.

Narciso C, Zartman J. 2018. Reverse-engineering organogenesis through feedback loops between model systems. Current Opinion in Biotechnology. 52:1–8.

Nebigil CG, Hickel P, Messaddeq N, Vonesch JL, Douchet MP, Monassier L, Gyorgy K, Matz R, Andriantsitohaina R, Manivet P et al. 2001. Ablation of Serotonin 5-HT2B Receptors in Mice Leads to Abnormal Cardiac Structure and Function. Circulation. 103:2973–2979.

Neumann CJ, Cohen SM. 1996. Distinct mitogenic and cell fate specification functions of wingless in different regions of the wing. Development (Cambridge, England). 122:1781–1789.

Nieto Gutierrez A, McDonald PH. 2018. GPCRs: Emerging anti-cancer drug targets. Cellular Signalling. 41:65–74.

Obata K, Shimo T, Okui T, Matsumoto K, Takada H, Takabatake K, Kunisada Y, Ibaragi S, Nagatsuka H, Sasaki A. 2016. Tachykinin Receptor 3 Distribution in Human Oral Squamous Cell Carcinoma. Anticancer Research. 36:6335–6341.

Obata K, Shimo T, Okui T, Matsumoto K, Takada H, Takabatake K, Kunisada Y, Ibaragi S, Yoshioka N, Kishimoto K et al. 2017. Role of Neurokinin 3 Receptor Signaling in Oral Squamous Cell Carcinoma. Anticancer Research. 37:6119–6123.

Padgett CL, Slesinger PA. 2010. GABAB Receptor Coupling to G-proteins and Ion Channels, In: Blackburn TP, Blackburn TP, editors, Advances in Pharmacology, Academic Press. volume 58 of GABAReceptor Pharmacology. pp. 123–147.

Pal K, Mukhopadhyay S. 2015. Primary cilium and sonic hedgehog signaling during neural tube patterning: Role of GPCRs and second messengers. Developmental Neurobiology. 75:337–348.

Pandey UB, Nichols CD. 2011. Human Disease Models in Drosophila melanogaster and the Role of the Fly in Therapeutic Drug Discovery. Pharmacological Reviews. 63:411–436.

Patel E, Kushwaha DS. 2020. Clustering Cloud Workloads: K-Means vs Gaussian Mixture Model. Procedia Computer Science. 171:158–167.

Patel M, Kawano T, Suzuki N, Hamakubo T, Karginov AV, Kozasa T. 2014. G*α*13/PDZ-RhoGEF/RhoA signaling is essential for gastrin-releasing peptide receptor-mediated colon cancer cell migration. Molecular Pharmacology. 86:252–262.

Pearson K. 1901. LIII. On lines and planes of closest fit to systems of points in space. The London, Edinburgh, and Dublin Philosophical Magazine and Journal of Science. 2:559–572.

Peel DJ, Milner MJ. 1990. The diversity of cell morphology in cloned cell lines derived from Drosophila imaginal discs. Roux’s archives of developmental biology. 198:479–482.

Perkins LA, Holderbaum L, Tao R, Hu Y, Sopko R, McCall K, Yang-Zhou D, Flockhart I, Binari R, Shim HS et al. 2015. The transgenic RNAi project at Harvard Medical School: resources and validation. Genetics. 201:843–852. Number: 3 Reporter: Genetics.

Perrimon N, Bonini NM, Dhillon P. 2016. Fruit flies on the front line: the translational impact of Drosophila. Disease Models & Mechanisms. 9:229–231.

Pinard A, Seddik R, Bettler B. 2010. GABAB Receptors: Physiological Functions and Mechanisms of Diversity, In: Blackburn TP, editor, Advances in Pharmacology, Academic Press. volume 58 of GABAReceptor Pharmacology. pp. 231–255.

R Core Team. 2022. R: A Language and Environment for Statistical Computing. R Foundation for Statistical Computing. Vienna, Austria.

Ralston A, Blair SS. 2005. Long-range Dpp signaling is regulated to restrict BMP signaling to a crossvein competent zone. Developmental Biology. 280:187–200.

Ray M, Lakhotia SC. 2019. Activated Ras/JNK driven Dilp8 in imaginal discs adversely affects organismal homeostasis during early pupal stage in Drosophila, a new checkpoint for development. Developmental Dynamics. 248:1211–1231.

Ren N, Zhu C, Lee H, Adler PN. 2005. Gene expression during Drosophila wing morphogenesis and differentiation. Genetics. 171:625–638.

Restrepo S, Zartman J, Basler K. 2014. Coordination of Patterning and Growth by the Morphogen DPP. Current Biology. 24:R245–R255. Number: 6.

Ries AS, Hermanns T, Poeck B, Strauss R. 2017. Serotonin modulates a depression-like state in Drosophila responsive to lithium treatment. Nature Communications. 8:15738.

Rotelli MD, Bolling AM, Killion AW, Weinberg AJ, Dixon MJ, Calvi BR. 2019. An RNAi Screen for Genes Required for Growth of Drosophila Wing Tissue. G3: Genes|Genomes|Genetics. 9:3087–3100.

Saad F, Hipfner DR. 2021. Extensive crosstalk of G protein-coupled receptors with the Hedgehog signalling pathway. Development. 148:dev189258.

Schulte G, Wright SC. 2018. Frizzleds as GPCRs – More Conventional Than We Thought! Trends in Pharmacological Sciences. 39:828–842.

Schwabe T, Bainton RJ, Fetter RD, Heberlein U, Gaul U. 2005. GPCR Signaling Is Required for Blood-Brain Barrier Formation in Drosophila. Cell. 123:133–144.

Sexton RE, Hachem AH, Assi AA, Bukhsh MA, Gorski DH, Speyer CL. 2018. Metabotropic glutamate receptor-1 regulates inflammation in triple negative breast cancer. Scientific Reports. 8:16008.

Shao Z, Masuho I, Tumber A, Maynes JT, Tavares E, Ali A, Hewson S, Schulze A, Kannu P, Martemyanov KA et al. 2021. Extended Phenotyping and Functional Validation Facilitate Diagnosis of a Complex Patient Harboring Genetic Variants in MCCC1 and GNB5 Causing Overlapping Phenotypes. Genes. 12:1352.

Skiba MA, Kruse AC. 2021. Autoantibodies as Endogenous Modulators of GPCR Signaling. Trends in Pharmacological Sciences. 42:135–150. Publisher: Elsevier.

Sobala LF, Adler PN. 2016. The Gene Expression Program for the Formation of Wing Cuticle in Drosophila. PLOS Genetics. 12:e1006100. Publisher: Public Library of Science.

Sommer C, Straehle C, Köthe U, Hamprecht FA. 2011. Ilastik: Interactive learning and segmentation toolkit. In:. pp. 230–233. Chicago, IL. ISSN: 1945-8452.

Sriram K, Insel PA. 2018. G Protein-Coupled Receptors as Targets for Approved Drugs: How Many Targets and How Many Drugs? Molecular Pharmacology. 93:251–258.

Strigini M, Cohen SM. 1999. Formation of morphogen gradients in the Drosophila wing. Seminars in Cell & Developmental Biology. 10:335–344.

Syrovatkina V, Alegre KO, Dey R, Huang XY. 2016. Regulation, Signaling and Physiological Functions of G-proteins. Journal of molecular biology. 428:3850–3868.

Szklarczyk D, Franceschini A, Kuhn M, Simonovic M, Roth A, Minguez P, Doerks T, Stark M, Muller J, Bork P et al. 2011. The STRING database in 2011: functional interaction networks of proteins, globally integrated and scored. Nucleic Acids Research. 39:D561–D568.

Szklarczyk D, Franceschini A, Wyder S, Forslund K, Heller D, Huerta-Cepas J, Simonovic M, Roth A, Santos A, Tsafou KP et al. 2015. STRING v10: protein–protein interaction networks, integrated over the tree of life. Nucleic Acids Research. 43:D447–D452.

Terunuma M. 2018. Diversity of structure and function of GABAB receptors: a complexity of GABAB-mediated signaling. Proceedings of the Japan Academy. Series B, Physical and Biological Sciences. 94:390–411.

The Gene Ontology Consortium. 2019. The Gene Ontology Resource: 20 years and still GOing strong. Nucleic Acids Research. 47:D330–D338.

Thurmond J, Goodman JL, Strelets VB, Attrill H, Gramates LS, Marygold SJ, Matthews BB, Millburn G, Antonazzo G, Trovisco V et al. 2019. FlyBase 2.0: the next generation. Nucleic Acids Research. 47:D759–D765.

Tiger M, Varnäs K, Okubo Y, Lundberg J. 2018. The 5-HT1B receptor - a potential target for antidepressant treatment. Psychopharmacology. 235:1317–1334.

Tuteja N. 2009. Signaling through G protein coupled receptors. Plant Signaling & Behavior. 4:942–947.

Wang D. 2018. The essential role of G protein-coupled receptor (GPCR) signaling in regulating T cell immunity. Immunopharmacology and Immunotoxicology. 40:187–192.

Wang L. 2005. Support Vector Machines: Theory and Applications. Springer Berlin, Heidelberg.

Werry TD, Wilkinson GF, Willars GB. 2003. Mechanisms of cross-talk between G-protein-coupled receptors resulting in enhanced release of intracellular Ca2+. The Biochemical Journal. 374:281–296.

Yang D, Zhou Q, Labroska V, Qin S, Darbalaei S, Wu Y, Yuliantie E, Xie L, Tao H, Cheng J et al. 2021a. G protein-coupled receptors: structure-and function-based drug discovery. Signal Transduction and Targeted Therapy. 6:1–27. Nature Publishing Group.

Yang MS, Lai CY, Lin CY. 2012. A robust EM clustering algorithm for Gaussian mixture models. Pattern Recognition. 45:3950–3961.

Yang Y, Zhang L, Yu J, Ma Z, Li M, Wang J, Hu P, Zou J, Liu X, Liu Y et al. 2021b. A Novel 5-HT1B Receptor Agonist of Herbal Compounds and One of the Therapeutic Uses for Alzheimer’s Disease. Frontiers in Pharmacology. 12.

Zhang Z, Xu Y. 2022. FZD7 accelerates hepatic metastases in pancreatic cancer by strengthening EMT and stemness associated with TGF-*β*/SMAD3 signaling. Molecular Medicine (Cambridge, Mass.). 28:82.

