## Supplementary Figures for "G protein-coupled Receptor Contributions to Wing Growth and Morphogenesis in *Drosophila melanogaster*"

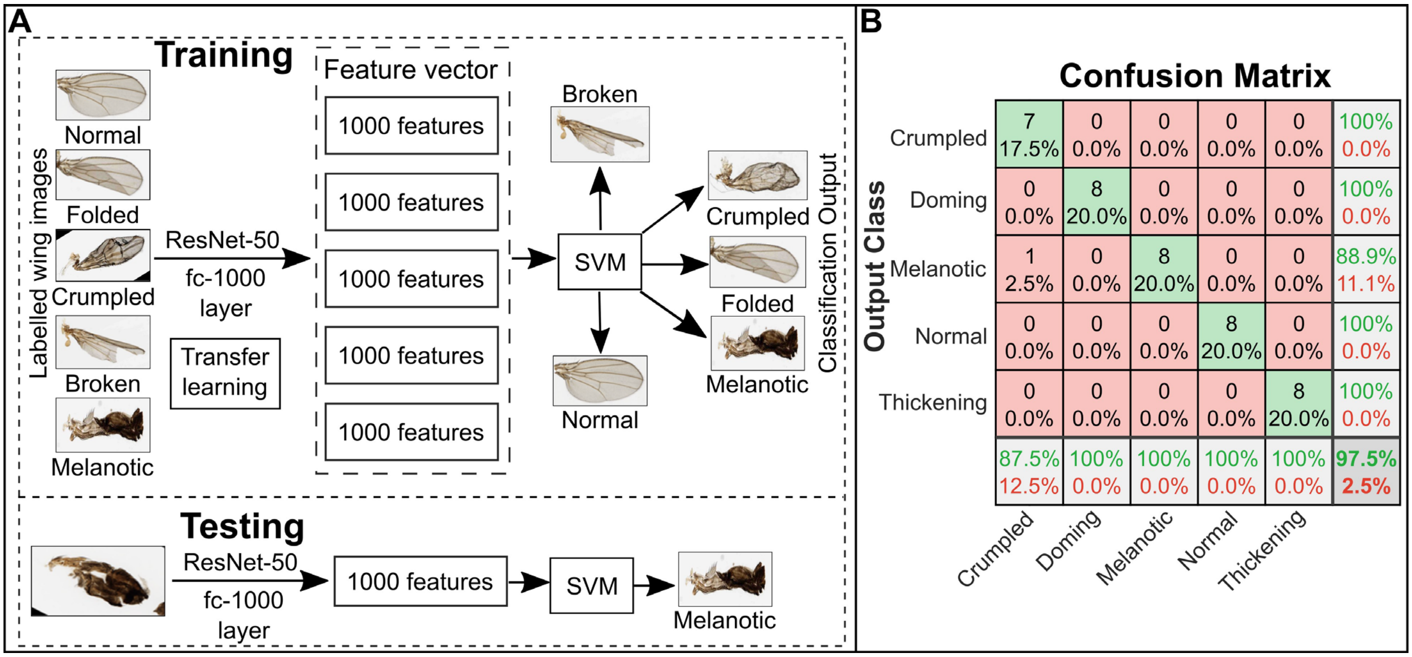


**SI Figure 1 | Machine learning-based pipeline fully automates the morphological analysis of wings. (A)** The fully connected (fc)-1000 layer of ResNet-50 was used to extract 1000 features from adult wing images in a training data set. The identified features of each wing were used to train a support-vector machine (SVM) classifier. The SVM classifier was then used to classify the wings into the different phenotypic classes observed after genetic knockdown of screened genes. **(B)** The confusion matrix of the accuracy for testing the SVM classifier is plotted. The confusion matrix demonstrates that the overall accuracy of the classifier is 97.5%. Accuracy of doming, melanotic, normal, and thickening classes is 100% with crumpled wings having an accuracy of 87.5%. The misclassified wing in the crumpled group was labeled as a melanotic wing, which is a reasonable misclassification because the two wing phenotypes are visually similar.

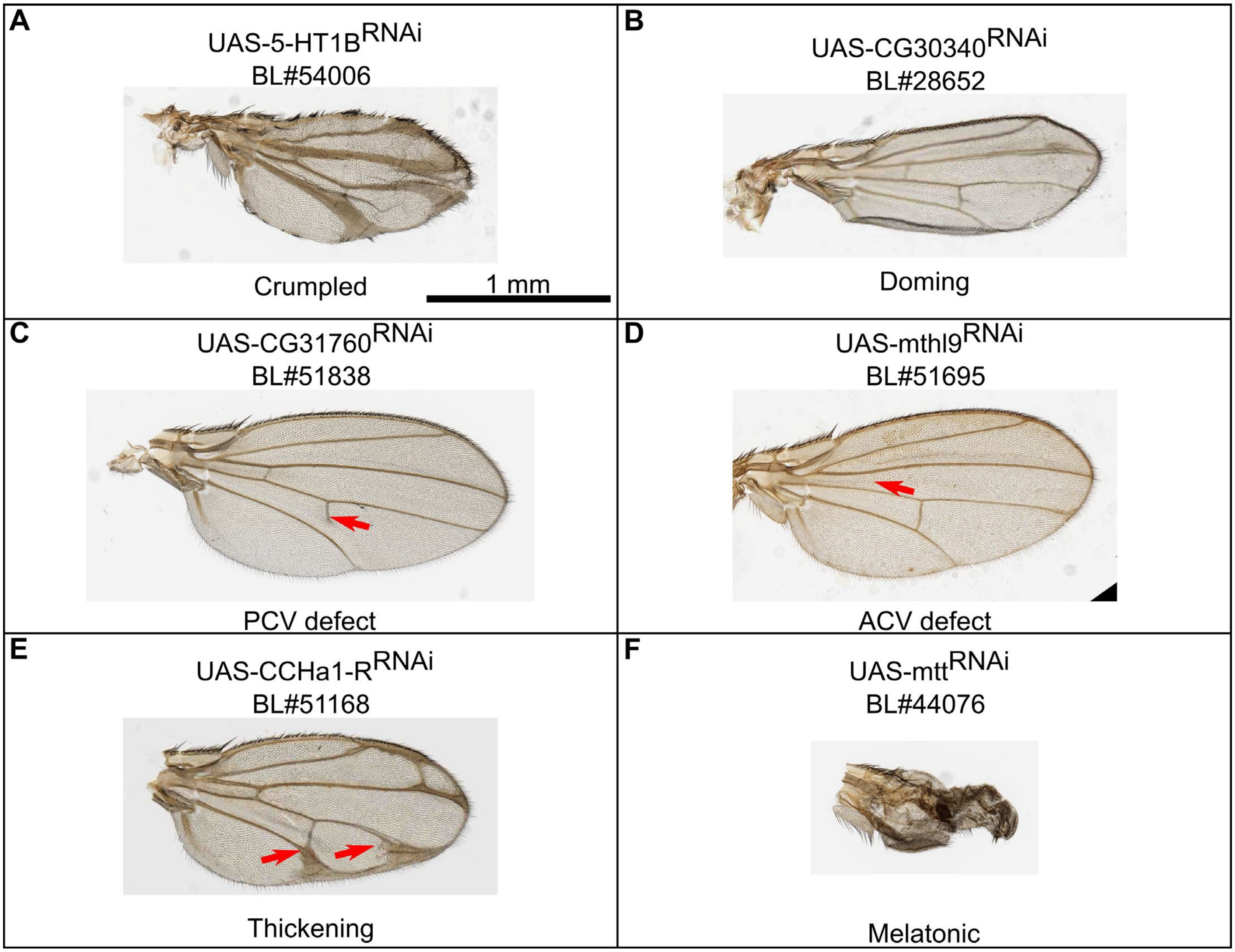


**SI Figure 2 | Various classes of severe phenotypes resulting from genetic knockdown experiments of GPCRs.** The MS1096-Gal4 (BL#25706) line was used as the basis for genetic crosses. Genetic knockdown progeny were generated by crossing the MS1096-Gal4 line to RNAi-based transgenic lines (UAS-Gene X$^{RNAi}$ (Perkins *et al.* 2015)). BL# is indicative of the Bloomington *Drosophila* Stock Center stock number for the genetic line used. Images are of adult male wings from F1 progeny resulting from MS1096-Gal4>UAS-Gene X$^{RNAi}$ crosses. The scale bar in panel **A** represents 1 mm and applies to all panels in the figure. **(A)** MS1096-Gal4>UAS-5-HT1B$^{RNAi}$ sample wing demonstrating the crumpled phenotype. **(B)** MS1096-Gal4>UAS-CG30340$^{RNAi}$ sample wing demonstrating the doming phenotype. **(C)** MS1096-Gal4>UAS-CG31760$^{RNAi}$ sample wing demonstrating a posterior crossvein (PCV) defect phenotype. **(D)** MS1096-Gal4>UAS-mthl9$^{RNAi}$ sample wing demonstrating an anterior crossvein (ACV) defect phenotype. **(E)** MS1096-Gal4>UAS- CCHa1-R$^{RNAi}$ sample wing demonstrating the thickening phenotype. **(F)** MS1096-Gal4>UAS-mtt$^{RNAi}$ sample wing demonstrating the melanotic phenotype.

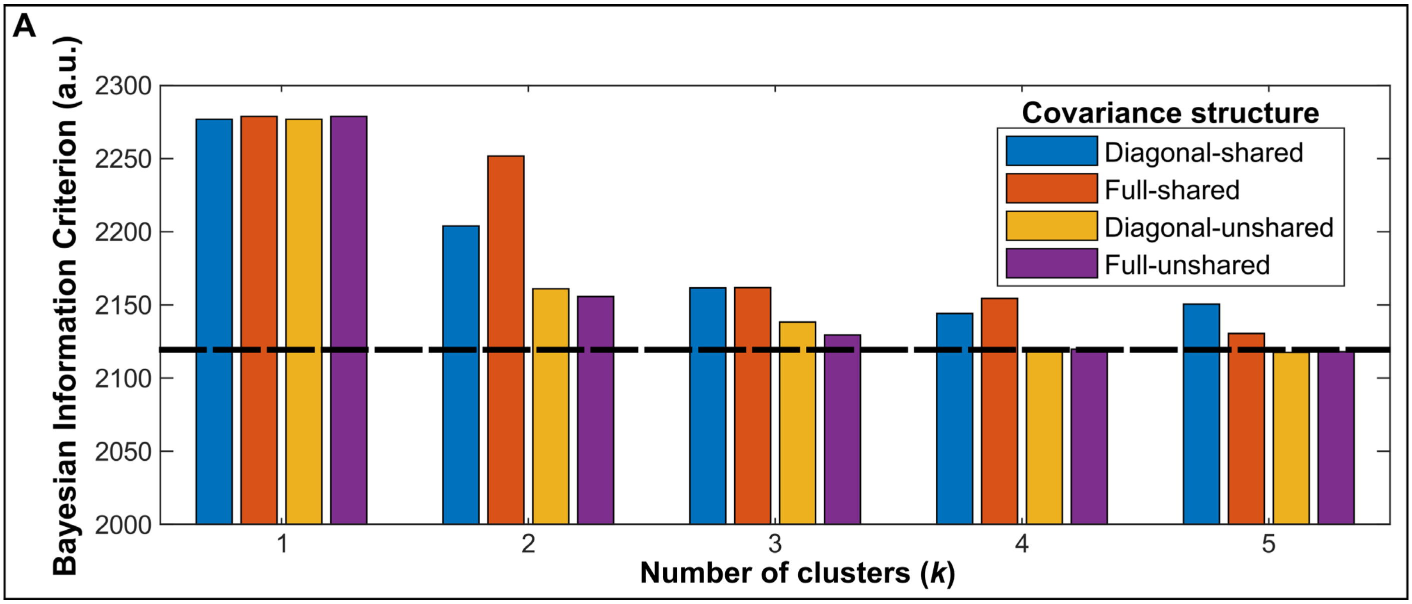


**SI Figure 3 | Bayesian information criterion identifies four clusters within the population of wing phenotypes.** For the Gaussian mixture model (GMM) clustering analysis, the number of clusters to fit to the GMM model was determined using the Bayesian information criterion (BIC). Optimizing the BIC for model selection criterion in conjunction with hierarchical clustering enables choosing the number of clusters to use in analysis according to the complexity present in the data set (Chen and Gopalakrishnan 1998). A lower BIC value indicates a better model fit. **(A)** GMMs of varying combinations of number of clusters ($k$) and covariance structures were fit to embedded features from the fully-connected layer of the ResNet-50 neural network. The features were first mapped to a two-dimensional space using principal component analysis. Plotted are the output BIC values for each combination of $k$ and covariance structure. The lowest BIC value occurs for $k = 4$ clusters with a covariance structure of full-unshared.

### SI Literature cited

Chen SS, Gopalakrishnan P. 1998. Clustering via the Bayesian information criterion with applications in speech recognition. In: . volume 2. pp. 645–648 vol.2. ISSN: 1520-6149.

Perkins LA, Holderbaum L, Tao R, Hu Y, Sopko R, McCall K, Yang-Zhou D, Flockhart I, Binari R, Shim HS *et al*. 2015. The transgenic RNAi project at Harvard Medical School: resources and validation. Genetics. 201:843–852. Number: 3 Reporter: Genetics
